# Time-resolved RNA-seq data reveal dynamic expressional behaviour of TALE target genes

**DOI:** 10.1101/2024.05.28.596194

**Authors:** Jan Grau, René Grove, Annett Erkes, Neele Unger, Marcel Göricke, Jens Boch

## Abstract

*Xanthomonas oryzae* bacteria infect rice (*Oryza sativa*) plants causing substantial harvest loss. During the infection, the bacteria translocate a collection of effector proteins into the host plant cells. This includes transcription activator-like effectors (TALEs) that act as transcriptional activators of plant genes. To understand the dynamics of TALE action during the infection, we collected RNA-seq data at different time points after the infection of rice plants with two representative pathogens of the two rice-pathogenic *Xanthomonas oryzae* ssp., namely *Xan-thomonas oryzae* pv. *oryzae* PXO83 and *Xanthomonas oryzae* pv. *oryzicola* BAI35. We observed that the induction of direct TALE target transcripts already starts 24h after the infection and is widely established after 36h, while 72h after the infection, secondary targets and downstream effects start to dominate the set of differentially expressed transcripts relative to control. Based on computational predictions of TALE targets combined with expression data, we established criteria that may help to identify direct TALE targets, which are related to expression dynamics but also shifted transcription start sites, and compiled a list of high-confidence TALE targets. Using genome-wide target predictions, we further discovered several non-coding and anti-sense transcripts that are likely induced by TALEs. Finally, we investigated different strategies to link putative secondary targets to transcription factors that are induced by TALEs during the infection.

## INTRODUCTION

Bacteria of the genus *Xanthomonas* are important plant pathogens that infect many crop plants, including rice, cotton, cabbage, citrus and barley (1). *Xanthomonas oryzae* ssp. infest rice (*Oryza sativa*) plants causing bacterial blight and leaf streak (1), which in consequence lead to massive yield losses. During the infection process, *Xanthomonas* bacteria translocate a collection of effector proteins into the host plant cells via a type III secretion system. In the plant cells, these effector proteins affect cellular processes and support bacterial virulence and proliferation, but may also trigger defence responses (2). One specific sub-type of such effector proteins are transcription activator-like effectors (TALEs) that bind to the host DNA and may specifically activate the expression of host plant genes, where a single *Xanthomonas* strain may harbour up to 28 TALEs (3). TALE proteins contain a nuclear localization signal, a modular DNA-binding domain that establishes target box specificity, and an activation domain that interacts with the host transcriptional machinery (4, 5). The DNA-binding domain is composed of up to 34 tandem repeats (median 19 repeats) and each repeat makes contact to one nucleotide of a contiguous DNA sequence. We refer to the DNA sequence bound by a TALE as *target box*. Each repeat contains approx. 34 amino acids, which fold into a helix-loop-helix structure. Amino acids 12 and 13 are located within the loop region and are responsible for the DNA specificity of the repeat (6). These amino acids are hyper-variable (repeat variable di-residue, RVD) and different combinations of amino acids lead to specific preferences for the DNA nucleotide bound by a repeat (6, 7). In addition, a cryptic repeat directly precedes the repeat array, which is also referred to as “position 0” and typically has a preference for nucleotide T. This modular structure of the DNA-binding domain allowed for the development of computational models that are capable of predicting the DNA specificity of a TALE based on the sequence of its RVDs (8–11). In previous benchmark studies, we could show that the PrediTALE model yields a favourable prediction performance (11). Hence, prediction of target boxes within promoters of putative target transcripts is performed using PrediTALE in this study. Regardless of the model used for predicting TALE target boxes, these predictions suffer from a substantial number of false positives (11), which can be moderately reduced by considering additional factors like chromatin structure and DNA methylation (12, 13). For this reason, target box predictions are typically complemented by expression data that serve as additional evidence of the activation of putative target transcripts (8, 11). Such expression data are recorded at varying time points after the infection, including 24 h or 48 h (11, 14– 16) and no consensus has been established, yet, which time point is suited best to identify direct TALE targets. There are several examples of transcription factors among the TALE target transcripts (16–18), which in turn may activate or repress the expression of further host genes as *secondary targets*, and the number of such secondary targets tends to increase with progressing infection. Previously, only few data sets have been published that specifically focus on the expression dynamics during the infection, including a collection of studies considering RNA-seq data (no replicates) of *Xoo* PXO99^*A*^ 12h, 24h, 36h, and 48h after the infection (19– 21).

With this study, we collect RNA-seq data of rice plants at different time points (24h, 36h, 48h and 72h) after the infection with two different TALE-carrying strains, namely *Xan-thomonas oryzae* pv. *oryzae* (*Xoo*) PXO83 and *Xanthomonas oryzae* pv. *oryzicola* (*Xoc*) BAI35. We chose these two strains, since they show clear infection symptoms, both have sequenced genomes including their TALome (3, 22), and represent the different infection strategies of *Xoo* and *Xoc. Xoo* bacteria colonize the xylem and spread through the vascular system, whereas *Xoc* bacteria infects parenchyma (23, 24). We consider differentially expressed transcripts at the different time points compared with mock inoculation as a control to investigate at which time point direct TALE effects are fully established and when secondary effects of TALEs become more prevalent. We further study to which extent time-resolved RNA-seq data are sufficient to identify secondary targets that are activated or repressed by plant transcription factors that are direct TALE targets themselves. Using promoter-centric and genome-wide predictions of TALE target boxes in combination with the expressional dynamics of putative target transcripts, we establish several criteria that may help to distinguish true TALE targets from false positive predictions and downstream effects. Genome-wide target box predictions with subsequent filtering using a refined and extended version of DerTALE (11) enable the identification of TALE-induced transcripts independent of existing gene annotations.

## METHODS

### Plant growth conditions and inoculation

*Oryza sativa* ssp. *japonica* cv. Nipponbare plants were grown under glasshouse conditions at 28°C during the 16h light period and 25°C (night) at 70% relative humidity. The second and third true leaves of 4-week-old plants were infiltrated with a needleless syringe and bacterial suspension of *Xoo* PXO83 or *Xoc* BAI35 with an OD_600_ of 0.5 in 10 mM MgCl_2_, or 10 mM MgCl_2_ as mock control.

### RNA extraction and sequencing

Leaves were inoculated in six spots in an area of approx. 5 cm. Per time point, two leaves of three rice plants were inoculated for each strain and control. Three replicate samples were taken 24h, 36h, 48h and 72h after inoculation, frozen in liquid nitrogen, and RNA was extracted using the RNeasy Mini Kit (QIAGEN GmbH/Hilden, Germany). Strand-specific libraries were sequenced on an Illumina NovaSeq instrument (Genewiz/Azenta) as 150 bp paired-end reads.

### Quality control

Quality control was performed using FASTQC version 0.11.9 and results were aggregated using MULTIQC v1.21. Per base and per sequence quality scores passed quality checks. Slight deviations in per base sequence content among the first 10 bases of reads were considered non-critical for subsequent mapping using STAR.

### Mapping reads to the rice genome

Reads were mapped to the rice genome (Nipponbare IRGSP-1.0 reference) obtained from http://rice.uga.edu/pub/data/Eukaryotic_Projects/o_sativa/annotation_dbs/pseudomolecules/version_7.0/all.dir/all.chrs.con using STAR (25) version 2.7.10b. The following non-standard parameters were set for the STAR mapping: `--alignIntronMax 20000` and `--alignMatesGapMax 20000` to prevent STAR from mapping very large introns that are implausible for rice transcripts; `--outSAMtype BAM SortedByCoordinate` to directly obtain sorted BAM files as STAR output.

### Quantification of transcript expression

Per-sample expression was quantified on the level of transcripts using FEATURECOUNTS version 2.0.6 based on the MSU7 annotation of the rice genome obtained from http://rice.uga.edu/pub/data/Eukaryotic_Projects/o_sativa/annotation_dbs/pseudomolecules/version_7.0/all.dir/all.gff3. Paired-end reads were counted on the level of fragments by setting parameters `-p --countReadPairs`, and on the level of exons, aggregated per transcript by setting `-t exon -g Parent`. Strand-specific libraries were indicated by parameter `-s 2`. To gain maximum sensitivity and concordance with manual inspection via IGV, we further set `-M -O --fraction` to also count multi-mapping reads, evenly distributed across multi-mapping sites, and for all overlapping exons.

### Identification of differentially expressed transcripts

Quantification results of FEATURECOUNTS were loaded into R (26) version 4.3.1 and further processed using the DESEQ2 package (27) version 1.40.2. RNA-seq samples of all four time points, for inoculation with *Xoo* PXO83, *Xoc* BAI35 and Mock control in all three replicates per condition were processed jointly using the `DESeqDataSetFromMatrix` function. The experiment design was specified as a two-factor design with one factor with three levels for the infection condition (*Xoo* PXO83, *Xoc* BAI35, Mock; Mock is base level) and one factor for the time point (24h, 36h, 48h, 72h; 24h is base level). The model was specified as `condition*time` to capture condition-specific and time-specific effects but also interactions between both, since different time-dependent dynamics between the different infection conditions are expected. Normalization of count values and determination of differentially expressed transcripts was performed using the `DESeq` function with parameter `test = “Wald”` to obtain contrast-specific p-values. Differentially expressed transcripts (DETs) were queried using the `results` function with parameters `name=“cond_<condition>_vs_Mock”` for the comparison of *Xoo* PXO83 and *Xoc* BAI35 infection with Mock inoculation at 24h (since this is the base level) and with parameters `contrast=list(c(“cond_<condition>_vs_Mock”, “cond<condition>.time<time>“)` for the remaining time points.

Resulting lists were further filtered by adjusted p-value (`padj` *<* 0.05) and by log2-fold change, where transcripts with a log2-fold change above 2 were counted as upregulated and transcripts with a log2-fold change below −2 were counted as down-regulated.

PCA-Plots of samples were generated based on variance-stabilized expression values (`vst` function of DESEQ2) using a modified version of the DESEQ2 `plotPCA` function. Alluvial plots of DETs were generated using the ggalluvial R-package version 0.12.5.

### Gene ontology term enrichment

GO term assignments of rice transcripts were obtained from http://rice.uga.edu/pub/data/Eukaryotic_Projects/o_sativa/annotation_dbs/pseudomolecules/version_7.0/all.dir/all.GOSlim_assignment. GO terms enriched in a set of transcripts were determined using a two-sided Fisher’s exact test (Hypergeometric test) using the R function `fisher.test`, correcting p-values for multiple testing using the `p.adjust` function using default parameters. Adjusted p-values and log-odds ratios were recorded for visualization.

### Prediction of TALE targets in promoters

A substantial fraction of transcripts in the MSU7 annotation lacks UTRs. Hence, mapped RNA-seq data and the MSU7 annotation were input into the ANNOTATIONFINALIZER module of GEMOMA (28, 29) to add UTR annotations to such transcripts. Afterwards, promoter sequences were extracted as 300 bp upstream of the transcription start site (TSS) until 200 bp downstream of the TSS or the start codon, whichever comes first, as proposed previously (8).

TALomes of *Xoo* PXO83 and *Xoc* BAI35 were extracted based on the AnnoTALE (3) predictions of TALE genes in the published genomes of *Xoo* PXO83 (3) and *Xoc* BAI35 (22), and corresponding RVD sequences were determined using AnnoTALE as listed in Supplementary Data S1 and S2, respectively.

PREDITALE (11) was used to predict target boxes of all TALEs present in the TALomes of *Xoo* PXO83 and *Xoc* BAI35, respectively, with default parameters and a significance threshold of 10^−3^. Resulting predictions were further filtered for the top 50 predictions per TALE, ranked by the best prediction score per gene.

### Genome-wide prediction and filtering using Der-TALEv2

PREDITALE was also applied for genome-wide predictions with a significance threshold of 10^−3^ and setting the parameter for the reverse strand-penalty to `r=0`. These predictions were converted to GFF format using a custom R script and added to the visualization of mapping results in the IGV genome browser (30, 31).

Genome-wide predictions were further used to search for differentially expressed regions in one condition (*Xoo* PXO83 or *Xoc* BAI35 infection) compared with Mock control. To this end, our previous approach DERTALE (11) was extended in a revised and extended version termed DERTALEV2. Briefly, DERTALE considers each predicted target box from the genome-wide predictions and searches for a contiguous region in its vicinity that shows elevated coverage (i.e. elevated numbers of mapped reads) in the respective condition relative to the control.

In DERTALEV2, we extend the previous method by several aspects. First, we explicitly consider the fragment orientation of the mapped reads in strand-specific libraries when computing coverage, which results in separate coverage tracks for each of the strands. Second, we only consider the coverage track in the orientation of the predicted target box. Although TALEs have been shown to activate the expression of down-stream transcripts, even if the TALE binds in reverse orientation relative to the orientation to the transcript (32), this is rather the exception. Hence, we decide for an increased specificity at the expense of a slightly reduced sensitivity in this case. Third, we modified the default value of the initial size of the region around the predicted target box that is searched for differential coverage to 500 bp, since activation of transcription at a larger distance is unlikely. Fourth, we reduced the default minimum length of the initial candidate region to 100 bp. If an initial region of sufficient coverage is found, this region is tested for further elongation in steps of 100 bp. Finally, and most importantly, we refine the criterion for calling a differentially expressed region. In the original DERTALE version, this has been based on the log2-fold change of read counts that has been computed from the average coverage in the infection condition and the average coverage in the control condition within the candidate region. In DERTALEV2, we extend this to a statistical analysis based on an F-test. Specifically, we consider two models, one base model using only the average log-coverage across infection *and* control replicates, and an alternative linear model using an additional factor for the condition (infection vs. control). The parameter for the condition-specific factor is determined by minimizing the squared error between the true, replicate-specific log-coverage values and the values predicted using the linear model. Afterwards, residues between the true, replicate-specific log-coverage values and the predictions of the base model (total average) are computed and, likewise, the residues between the true, replicate-specific log-coverage values are computed. We denote the sum of the squared residues of the base model as *SSR*_0_ and the sum of the squared residues of the linear model as *SSR*_1_.

The F-statistic is then defined as

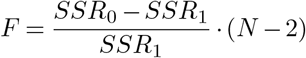

where *N* denotes the number of replicates (6 in our case) across infection condition and control, the linear model has 2 parameters (intercept and coefficient for one two-level factor) and the base model has one parameter (intercept).

Under the null hypothesis that the predictions of the linear model are as accurate as the predictions of the base model (besides random effects), this test statistic is F-distributed with *d*_1_ =1 and *d*_2_ = *N*−2 degrees of freedom. Under this null hypothesis, we may, hence, compute critical values of the F-statistic for a given significance level and p-values from the thus parameterized F-distribution. Here, we apply a significance level of *α* = 0.05 and report differentially expressed regions with a p-value below the significance level.

For convenience, DERTALEV2 outputs predicted TALE target boxes filtered by the presence of a differentially expressed region, and the detected differentially expressed region in GFF format.

For the time-resolved data considered in this study, we apply DERTALEV2 independently for each infection condition (*Xoo* PXO83 and *Xoc* BAI35) and each time point. Detected differentially expressed regions are further filtered by the presence of differentially expressed regions associated with the same TALE target box at multiple time points.

### Processing of ChIP-seq data for OsTFX1

ChIP-seq data of an OsTFX1 overexpression line are publicly available (33). We considered these data on three levels of processing. First, we obtained the list of ChIP-associated genes from Supplementary Data 2 of the original publication. Second, we obtained ChIP-seq peaks of two replicates from ChIP-Hub (34) that are based on the same raw data but have been processed using the unified pipeline of ChIP-Hub. Third, we downloaded the raw reads, which are available from the NCBI sequence read archive under accession PRJNA450934. We mapped these reads to the rice genome using bowtie2 (35) version 2.3.5.1 with parameter `--local`. We then called peaks using MACS (36) version for sample SRR7039127 against control SRR7039129, and for sample SRR7039128 against control SRR7039130. We further established the association between peaks and transcripts by applying the `bedtools closest` command of BED-TOOLS (37) to the respective peak file and the rice genome annotation.

For the ChIP-seq peaks obtained from ChIP-Hub and called via MACS, we considered the union of peaks for down-stream analyses.

### Motif-based prediction of TF binding sites

Binding motifs of LOC_Os01g46800.1 (M07399_2.00), LOC_Os01g61080.1 (M07391_2.00), LOC_Os01g63510.1 (M06905_2.00) were obtained from CIS-BP (38). A binding motif of the *A. thaliana* homolog AT5G38800.1 of LOC_Os09g29820.1 was obtained from PlantTFDB (https://planttfdb.gao-lab.org/motif/Ath/AT5G38800.meme) as it matches the consensus of the PBM-based motifs in CIS-BP (M01826_2.00, M01827_2.00) but is longer and more specific. In addition, this motif is highly similar to the motif obtained by applying our motif discovery tool Dimont (39) to the OsTFX1 ChIP-seq peaks. All motifs with the corresponding sequence logos are listed in Supplementary Table C.

To find significant binding sites for these motifs, Fimo (40) was applied using the previously extracted promoter sequences as input. The significance threshold of Fimo was set to 10^−5^. The output files of Fimo (`fimo.tsv`) were considered in downstream analyses.

### Generation of co-expression networks

RNA-seq data of 785 samples of rice were downloaded from ENA. The corresponding accessions are listed in Supplementary Table E. Expression of annotated rice transcripts according to http://rice.uga.edu/pub/data/Eukaryotic_Projects/o_sativa/annotation_dbs/pseudomolecules/version_7.0/all.dir/all.cdna was quantified using KALLISTO (41) version 0.48.0. Single-end reads were processed with additional parameters `--single -l 200 -s 80`. Quantification results (output file `abundance.h5`)) were loaded into R using the `tximport` package and normalized using DESEQ2. Variance stabilized abundance values were determined using the `vst` function of DESEQ2. These served as input of the re-implementation of the GENIE3 method (42) within the ARBORETO re-implementation that is part of the SCENIC framework (43). Specifically, this expression matrix was loaded into python using the `read_csv` function of PANDAS and the co-expression network was determined from that matrix using the `genie3` function of ARBORETO. As putative TFs (argument `tf_names`), all rice transcripts with GO term “GO:0003700” according to the rice GO annotation (cf.) or a binding motif in CIS-BP were considered. The resulting network was saved as a TSV file using the `to_csv` method of the `genie3` object. This table contains one row for each edge, including the corresponding edge weight, which can be used for ranking and filtering TF targets.

## RESULTS/DISCUSSION

To characterize how TALEs from *Xoo* PXO83 and *Xoc* BAI35 trigger the induction of rice transcripts over the time course of the initial infection, we sampled RNA at 24h, 36h, 48h, and 72h post inoculation and quantified the expression of rice transcripts. Differentially expressed genes at each time point were determined relative to Mock inoculation as a control. In the following, we analyze differentially expressed transcripts for their expression dynamics, combine these with computational predictions to identify high-confidence, direct TALE targets, and investigate putative secondary targets.

### Overview of sequencing libraries

Rice leafs were collected 24h, 36h, 48h and 72h after *Xoo* PXO83 infection, *Xoc* BAI35 infection and Mock inoculation, RNA was extracted and sequenced in paired-end, strand specific libraries. After initial quality control using FASTQC, reads were mapped to the rice genome using STAR (25) and assigned to individual transcripts using fea-tureCounts (44). The number of reads in all libraries was sufficient for reliable estimation of transcript abundance, and mapping rates were reasonable (Supplementary Table A). Quantification results were further processed using the DESEQ2 (27) R-package.

Figure 1 shows a 2D PCA (principle component analysis) embedding of the per-sample expression vectors across all libraries. In general, we find good concordance of biological replicates except for one replicate 72h after *Xoc* BAI35 infection that is similar to the respective 24h replicates, which will reduce statistical power when comparing *Xoc* BAI35 infection vs. Mock inoculation at 72h. Notably, all replicates taken at 36h behave markedly different than the other replicates as indicated by their spread across PC1. This is likely an effect of the circadian rhythm of the rice plants, but does not affect comparisons between infection and Mock inoculation, in general, since comparisons are always performed within identical time points. Supplementary Figure S1 shows the PCA-plot when excluding all 36h replicates, which avoids this effect. Here, we observe that all replicates at 24h start similarly, while samples after *Xoo* PXO83 and *Xoc* BAI35 infection progress similarly along PC1, but in different directions along PC2. This gives a first indication that the general progression of infection is similar for *Xoo* PXO83 and *Xoc* BAI35, but subsets of transcripts that respond to the infection differ between *Xoo* PXO83 and *Xoc* BAI35.

**Fig. 1.**
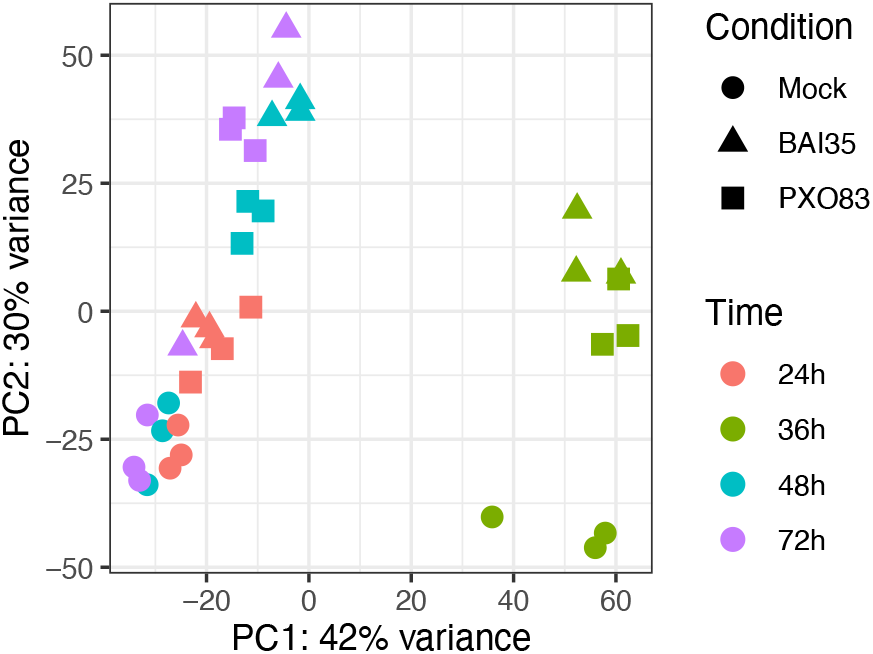
PCA plot of the quantification results of all replicates, where shape indicates condition (infection with *Xoo* PXO83 or *Xoc* BAI35; Mock inoculation) and colour indicates time point.

### Overview of differentially expressed transcripts

To identify TALE-dependently induced rice transcripts, we searched for transcripts that are differentially expressed in comparison to Mock inoculation and contain a predicted TALE target box in their promoter.

Differentially expressed transcripts were determined using DESeq2 (Wald test, adjusted p-value *<* 0.05, absolute log2-fold change *>* 2) testing *Xoo* PXO83 and *Xoc* BAI35 infection against Mock inoculation at 24h, 36h, 48h and 72h, respectively. In addition, TALE targets were predicted using PrediTALE (11) for all TALEs present in the TALomes of *Xoo* PXO83 (3) and *Xoc* BAI35 (22) considering putative promoter sequences from 300 bp upstream of the annotated transcription start site (TSS) to 200 bp downstream of the TSS or the start codon, as proposed previously (8). Here, we consider the top 50 predictions per TALE as in previous benchmark studies (11).

Figure 2 A and B show alluvial plots of up-regulated, differentially expressed transcripts (DETs) for *Xoo* PXO83 and *Xoc* BAI35, respectively, over the time course. Per time point, transcripts are assigned to the categories “DET and TALE target” if a transcript is considered differentially expressed and yields a predicted TALE target box in its promoter, “only DET” and “only TALE target” if only one of the conditions holds, or “Neither” if it is neither differentially expressed nor a predicted TALE target. We observe that the number of DETs is rather low at 24h, increases substantially until 36h, stays at a similar level at 48h, and again increases at 72h. The number of direct TALE targets (“DET and TALE target”) is generally low, while the majority of DETs is likely due to the general response of the host plant to the infection – especially at early time points – or due to potential secondary effects of early TALE targets – especially at 72h. We also observe that direct TALE targets that are established at earlier time points typically remain direct TALE targets at later time points, where transcripts switch from category “only TALE target” to “DET and TALE target”. The increase of direct TALE targets is especially pronounced going from the 24h time point to the 36h time point.

**Fig. 2.**
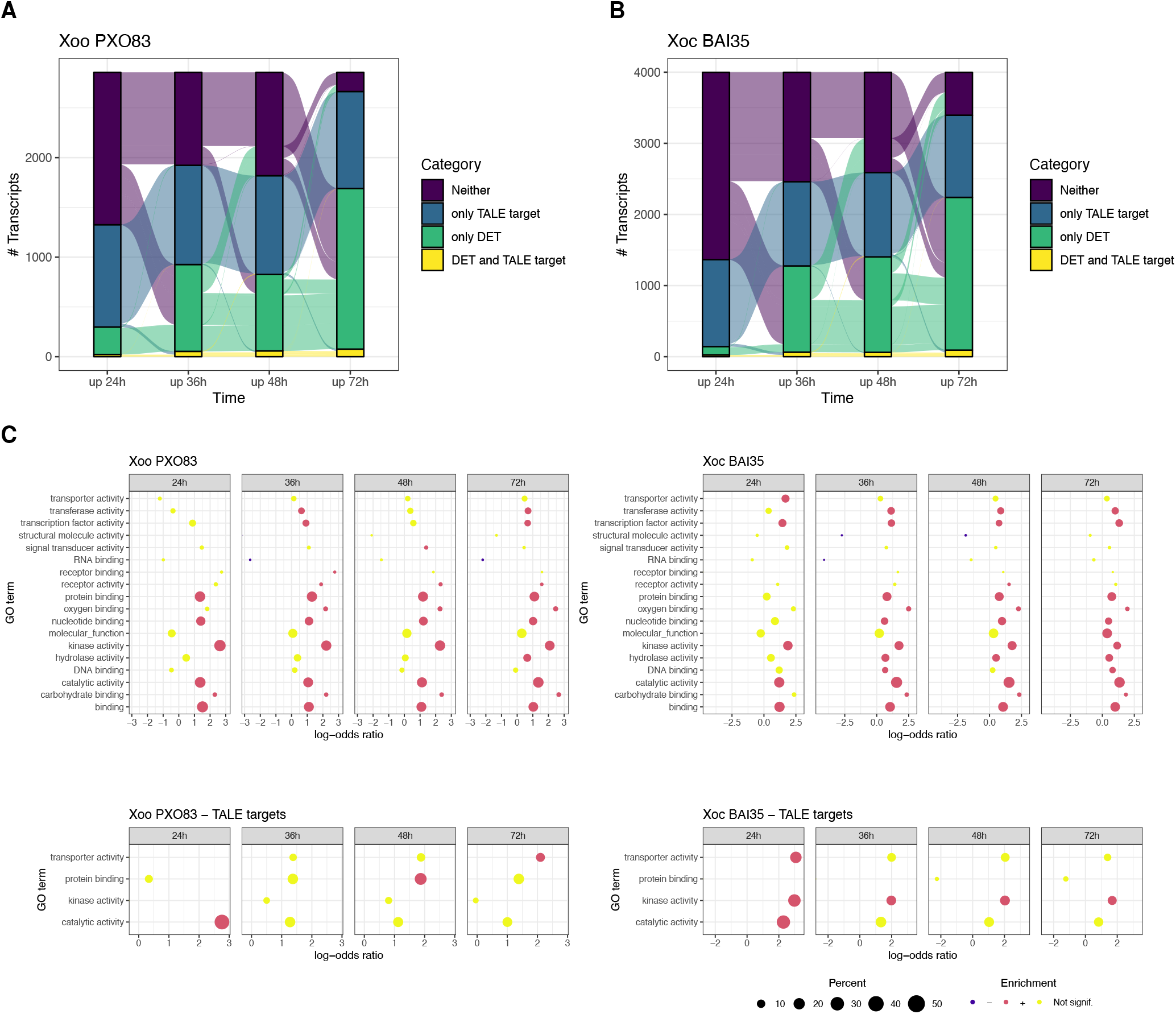
Alluvial plots of differentially expressed transcripts after *Xoo* PXO83 (A) and *Xoc* BAI35 (B) infection at the different time points. For each time point, transcripts are classified as “DET and TALE target” if they are differentially expressed and contain a predicted TALE target box in their promoter, “only TALE target” or “only DET” if only one of these conditions holds, or “Neither” otherwise. Here, we exclude transcripts that would be assigned to the “Neither” class at all time points. (C) Gene ontology (molecular function) terms enriched among the DETs at each of the time points for *Xoo* PXO83 (left) and *Xoc* BAI35 (right) and separately for the respective TALE targets (bottom).

Down-regulated DETs also exist (Supplementary Figure S2), but are considerably lower in number than up-regulated DETs and are almost devoid of predicted TALE targets, as can be expected due to the function of TALEs as transcriptional activators. This observation also increases our confidence that predicted TALE targets among up-regulated DETs are enriched with true, functional TALE targets as opposed to false-positive predictions.

From this overview evaluation of the transcriptional response to the *Xanthomonas* infection, we may already conclude that the effect on direct TALE targets already begins at 24h and is widely established at 36h, whereas DETs that appear later during the time course are mostly due to secondary effects.

In Figure 2C, we further consider enriched GO terms (molecular function) among all DETs at individual time points and specifically among the predicted direct TALE targets. Here, we find several general terms (e.g., “binding”, “catalytic activity”) enriched for both pathovars across all time points. Among the more specific terms, “transporter activity”’ is enriched at 24h after *Xoc* BAI35 infection (also for TALE targets), “transferase activity” is enriched at multiple time points for both pathovars, “transcription factor activity” is enriched at all time points for *Xoc* BAI35, and at 36h and 72h for *Xoo* PXO83, and “protein binding” and “nucleotide binding” are enriched at all time points except 24h after *Xoc* BAI35 infection. This picture is reflected for the enriched terms of the “biological process” ontology (Supplementary Figure S3) with significant enrichment of general terms like “metabolic process” but also more specific terms like “signal transduction”, “response to stress” or “response to biotic stimulus”. For the “cellular component” ontology (Supplementary Figure S4), we find “cell wall” and “plasma membrane” enriched, the latter also among the direct TALE targets of *Xoc* BAI35.

In addition, Figure 3 shows upset plots of the sets of DETs at the different time points and the percentage of predicted TALE targets per intersection set. Here, we observe a substantial number of direct TALE targets within the intersections across all time points and for the intersection of the 36h to 72h time points, which supports our previous conclusion that the TALE-induced up-regulation of transcripts is widely established at the 36h time point. Some direct TALE targets may also be found among the DETs specific for 72h, but their percentage is rather low and we consider these mostly false-positive predictions due to the large number of DETs at either 72h time point. We further consider joint upset plots of DETs after *Xoo* PXO83 and *Xoc* BAI35 infection in Supplementary Figure S5. Here, we distinguish between TALE targets that are predicted for both pathovars and those predicted exclusively for one of the pathovars. Among the intersection of DETs present across all time points and both pathovars, we find one common TALE target, LOC_Os01g40290, that is strongly induced (Supplementary Table D) after *Xoo* PXO83 infection (log2-fold change between 5.4 and 7.1) and after *Xoc* BAI35 infection (log2-fold change between 9.2 and 11.6) and is a predicted target of Ta-lAA5 (rank 1) of *Xoo* PXO83 and of TalCJ5 (rank 1) of *Xoc* BAI35. This gene does not have an annotated function, but the common, strong and early induction of its expression after infection with both pathovars indicates an important role in the infection process. The previously reported convergent targets of *Xoo* and *Xoc OsDOX-1* (LOC_Os03g03034.1) and *OsDOX-2* (LOC_Os04g49194.1) (16) are targeted by TalAQ3 (*Xoo* PXO83) and TalBL13 (*Xoc* BAI35), respectively, and show up-regulation in our experiments. As further examples of convergent TALE targets between *Xoo* and *Xoc*, OsLsi1 (LOC_Os02g51110.1; TalAL in *Xoo* and TalAV in *Xoc*) and OsFBX109 (LOC_Os03g51760.1; TalAD in *Xoo* and *Xoc*) have been reported (16), but do not as overlapping targets in our results, since *Xoo* PXO83 lacks a TalAL-class TYLE and *Xoc* BAI35 lacks a TalAD-class TALE. However, OsLsi1 is up-regulated after *Xoc* BAI35 infection and OsFBX109 is up-regulated after *Xoo* PXO83 infection (Supplementary Table D).

**Fig. 3.**
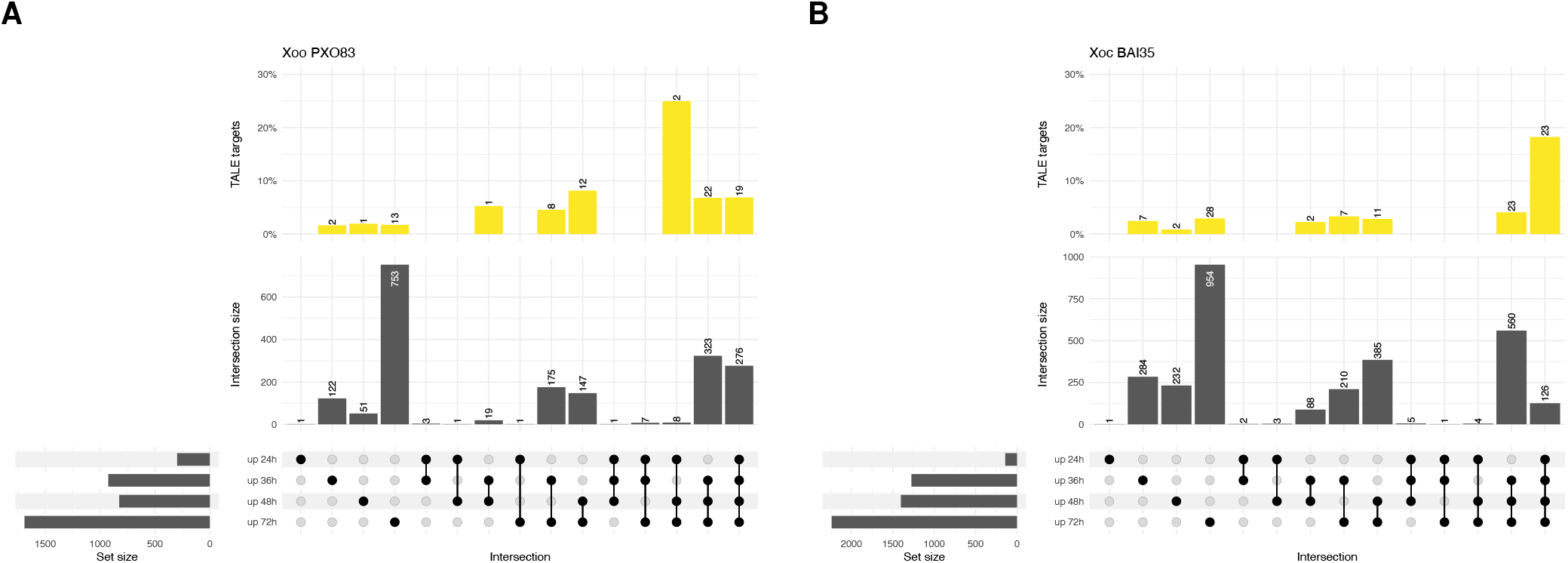
Upset plots of the sets of up-regulated transcripts at the individual time points after *Xoo* PXO83 (A) and *Xoc* BAI35 (B) infection. Subsets that are specific for one of the input seqs (left) are indicated by individual filled dots in the intersection matrix, while intersections of two or more sets are indicated by filled dots connected by lines. Intersection sizes are visualized by a bar plot above the intersection matrix. For each intersection, we further visualize the percentage of predicted TALE targets as barplots and provide the absolute numbers of predicted TALE targets above the respective bar.

We further find that the intersections between all four time points and between 36h to 72h exclusively for *Xoo* PXO83 is clearly enriched for predicted targets of *Xoo* PXO83 TALEs, while the analogous observation holds true for the respective intersections of *Xoc* BAI35. Again, this indicates that the predicted TALE targets among the respective DETs are true TALE targets for the most part.

According to this interpretation, we analyze the expressional behaviour of DETs present from 36h to 72h in more detail in the following.

### Clustering of DETs

We cluster transcripts by their expressional behaviour across all treatments and time points using the `degPatterns` function of the DEGREPORT R-package (45). Figure 4 visualizes the clustering result for up-regulated DETs in the intersection of time points 36h to 72h for *Xoo* PXO83 and *Xoc* BAI35. For *Xoo* PXO83, we observe one cluster (Group 3) comprising transcripts that are up-regulated exclusively in the *Xoo* PXO83 experiments but not after *Xoc* BAI35 infection or Mock inoculation. For Group 1 DETs, some response can also be observed after *Xoc* BAI35 infection, while for Group2 the response after infection with both pathovars is largely similar. All three clusters contain putative TALE targets, where the strongest enrichment (37.5%) is in Group 3. Genes in Group 2 and 3 show increasing expression over time after *Xoo* PXO83 infection, whereas for Group 1, expression initially increases but stays on similar levels from 36h to 72h. For *Xoc* BAI35, the general picture is similar with one cluster (Group 1) with response exclusively after *Xoc* BAI35 infection, one cluster (Group 3) with a slight response after *Xoo* PXO83, and one cluster (Group 2) with similar responses for both pathovars. However, the number of DETs that are exclusively up-regulated after the infection is larger for *Xoc* BAI35 than for *Xoo* PXO83. Also, the expressional dynamics after *Xoc* BAI35 infection differ, where Group 1 and 2 DETs reach their final level at 36h, already, whereas Group 3 DETs show increasing expression up to 48h.

**Fig. 4.**
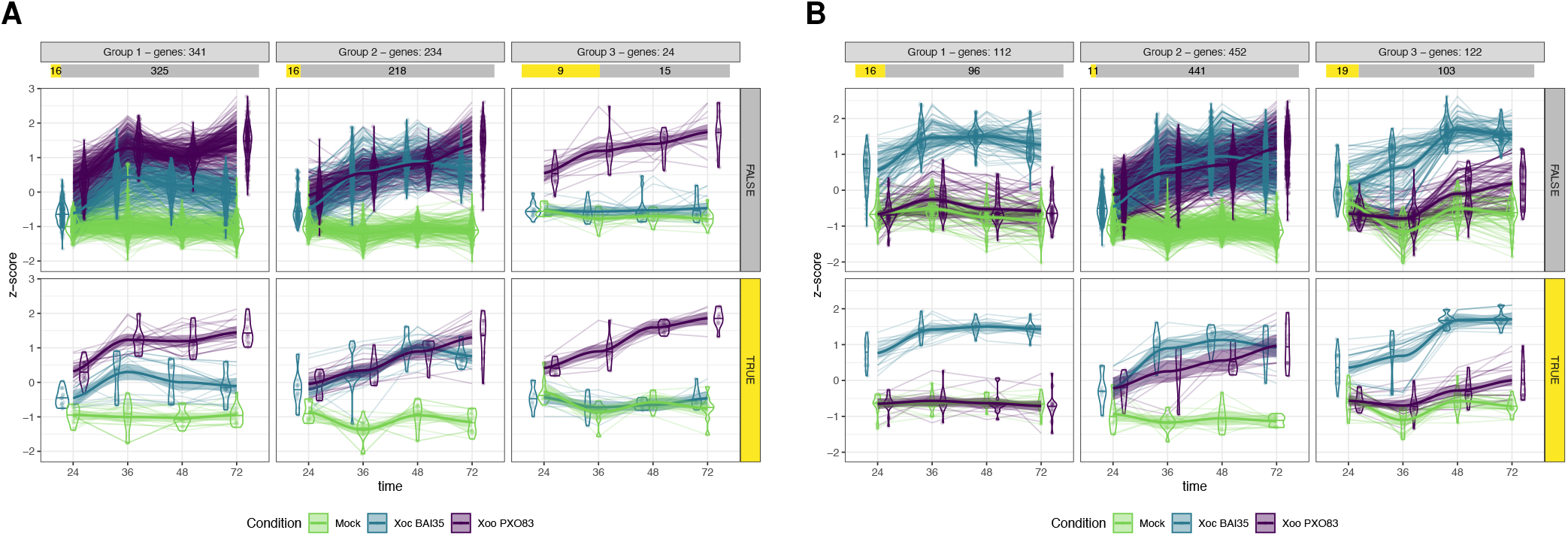
Clustering result of the expression profiles of all DETs that are consistently up-regulated from 36h to 72h after *Xoo* PXO83 (A) and *Xoc* BAI35 (B) infection. For each cluster, we separately plot expression profiles of putative TALE targets (bottom) and all remaining DETs (top), and visualize the proportions of TALE targets and other DETs below the cluster title bar. Expression profiles across time points are coloured based on the respective condition (Mock inoculation; *Xoo* PXO83, *Xoc* BAI35 infection).

Based on the bulk RNA-seq data of fixed amounts of input plant material considered in this study, explanation of (observed) increasing expression levels may be two-fold. On the one hand, continuous transcriptional activation by TALEs may lead to an accumulation of the respective transcripts in infected cells. On the other hand, the progression of infection may result in a larger proportion of plant cells into which effector proteins have been translocated and, hence, a larger proportion of plant cells with up-regulated TALE targets.

### High-confidence TALE targets

We list putative TALE targets that show significant and consistent up-regulation from 24h or 36h to 72h in Supplementary Table B. Here, we find several target genes that are up-regulated early and very strongly compared with Mock inoculation as listed in Table 1. For the two most extreme examples, namely LOC_Os01g63510.1 in *Xoo* PXO83 and LOC_Os06g46500.1 in *Xoc* BAI35, we provide snapshots of the IGV genome browser for the respective genomic regions in Supplementary Figures S6 and S7. Both examples are representative of many putative TALE targets with strong up-regulation listed in Table 1. The expression of LOC_Os01g63510.1 is likely induced by TalCA1 and shows expression levels (visible as number of mapped reads in the IGV snapshot) increasing over time. This transcript is (almost) absent in Mock inoculation but also after *Xoc* BAI35 infection, which gives additional indication of TALE-dependent up-regulation as opposed to general plant response to the infection. By contrast, LOC_Os06g46500.1 shows base-level expression in Mock inoculation (35 reads on average) and after *Xoo* PXO83 infection (15 reads on average), but is up-regulated to extreme levels (25, 786 reads on average; maximum of 71, 198 reads at 72h) after *Xoc* BAI35 infection, likely due to transcriptional activation by TalBF17. This demonstrates that following host transcriptomics over time during the onset of the infection enables us to make more solid predictions about TALE targets and to identify new ones.

**Table 1.** High-confidence TALE targets with early and strong up-regulation. Log2-fold changes (columns with prefix “lfc”) are provided for the infection with the respective strain relative to Mock inoculation.

| Strain | Transcript | lfc 24h | lfc 36h | lfc 48h | lfc 72h | TALE | Annotation |
| --- | --- | --- | --- | --- | --- | --- | --- |
| *Xoo* PXO83 | LOC_Os01g40290.1 | 5.41 | 6.40 | 7.05 | 7.11 | TalAA5 | expressed protein |
| *Xoo* PXO83 | LOC_Os01g43280.1 | 5.11 | 7.69 | 7.07 | 8.48 | TalBJ2 | hydroquinone glucosyltransferase |
| *Xoo* PXO83 | LOC_Os01g63510.1 | 6.72 | 7.58 | 9.25 | 10.23 | TalCA1 | homeobox domain |
| *Xoo* PXO83 | LOC_Os04g05050.1 | 4.01 | 4.59 | 5.22 | 6.93 | TalAB5 | pectate lyase precursor |
| *Xoo* PXO83 | LOC_Os04g19960.1 | 3.55 | 9.29 | 8.14 | 9.87 | TalAC5 | retrotransposon protein |
| *Xoo* PXO83 | LOC_Os07g06970.1 | 3.75 | 2.81 | 5.49 | 5.85 | TalAP3 | HEN1 |
| *Xoo* PXO83 | LOC_Os07g23410.1 | 3.24 | 6.91 | 8.14 | 9.78 | TalAA5 | fatty acid desaturase |
| *Xoo* PXO83 | LOC_Os09g29820.1 | 3.95 | 6.93 | 4.61 | 4.61 | TalAR3 | OsTFX1 |
| *Xoo* PXO83 | LOC_Os12g24320.1 | 4.59 | 4.39 | 5.63 | 6.94 | TalAF4 | ATPase 3 |
| *Xoc* BAI35 | LOC_Os01g05880.1 | 7.23 | 8.46 | 7.59 | 8.05 | TalCH5 | OsFBX1 |
| *Xoc* BAI35 | LOC_Os01g40290.1 | 9.26 | 10.35 | 11.64 | 10.64 | TalCJ5 | expressed protein |
| *Xoc* BAI35 | LOC_Os01g52130.1 | 8.82 | 13.12 | 10.4 | 10.96 | TalBF17 | sulfate transporter |
| *Xoc* BAI35 | LOC_Os01g16470.1 | 6.83 | 9.8 | 11.16 | 11.39 | TalCJ5 | phosphatidylinositol kinase |
| *Xoc* BAI35 | LOC_Os04g49194.1 | 8.30 | 11.50 | 10.92 | 12.67 | TalBL13 | OsDOX-2 |
| *Xoc* BAI35 | LOC_Os06g22290.1 | 4.81 | 9.96 | 7.64 | 9.34 | TalBF17 | legume lectins beta domain |
| *Xoc* BAI35 | LOC_Os06g46500.1 | 6.80 | 9.52 | 10.91 | 13.07 | TalBF17 | monocopper oxidase |

### Shifted transcription start sites

We further investigate putative TALE targets listed in Supplementary Table B for representative patterns of transcriptional activation by TALEs based on IGV snapshots. One pattern that occurs frequently are shifted transcription start sites (TSSs). Due to limitation of the UTR annotation present in the MSU7 genome annotation, these can be reliably identified only if base levels of expression are present in Mock inoculation. One example of a target gene with shifted TSS is given in Figure 5. LOC_Os09g29820.1 (OsTFX1) is a well known target of TalAR-class TALEs (16, 17, 46) and in this case is a predicted target of TalAR3 from *Xoo* PXO83. In Mock inoculation and after *Xoc* BAI35 infection, OsTFX1 is present at base levels, while expression is substantially increased after *Xoo* PXO83 infection already at 24h and expression further increases at later time points. Base level expression from the natural TSS is still present after *Xoo* PXO83 infection, but increased expression starts from a TSS approx. 40 bp down-stream (in transcript direction) from the natural TSS. A similar example for *Xoc* BAI35 is given in Supplementary Figure S8, where the TSS of LOC_Os06g37080.1 is shifted by approx. 40 bp after transcriptional activation by TalBC14.

**Fig. 5.**
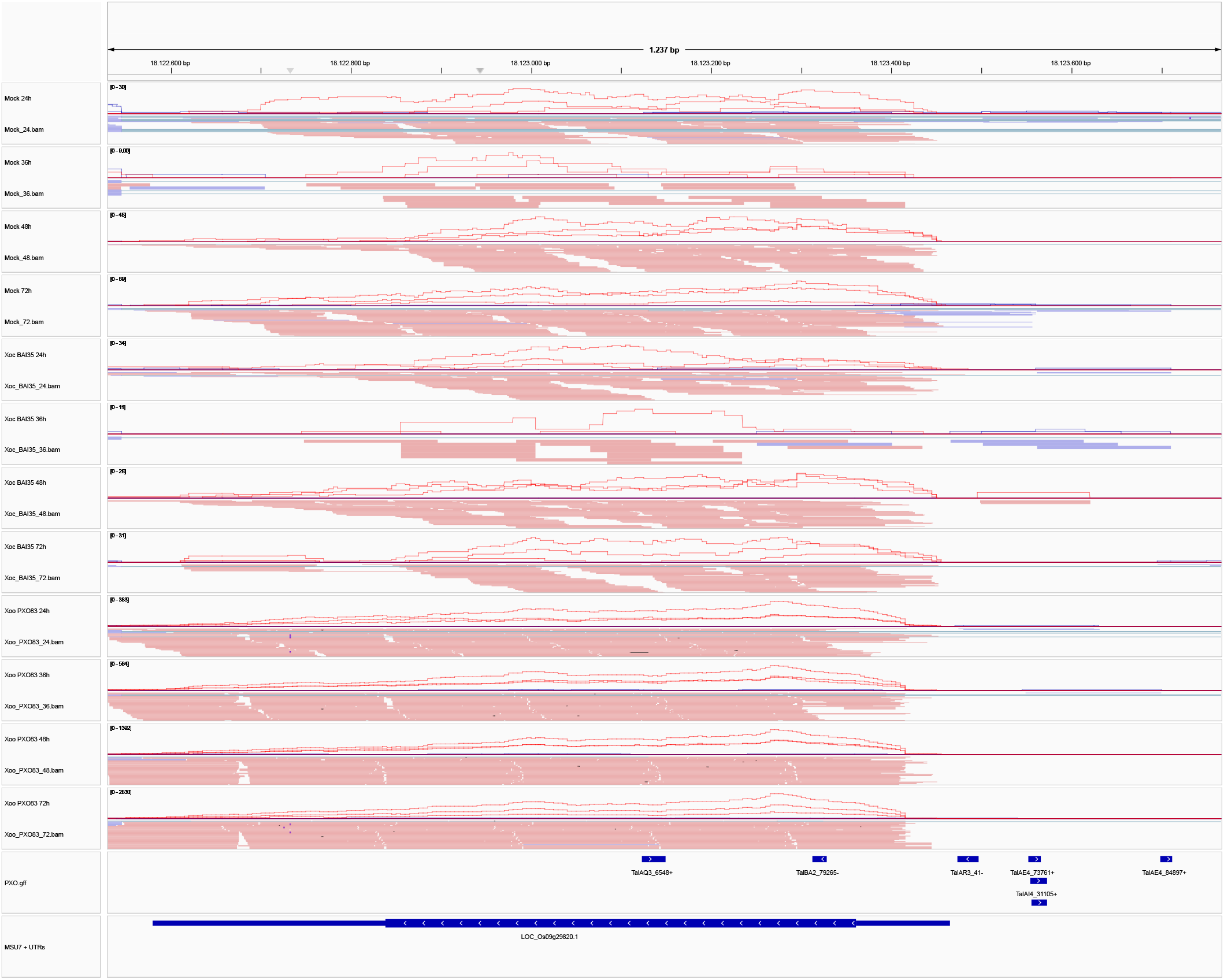
IGV snapshot of the genomic locus of LOC_Os09g29820.1 (OsTFX1). In each panel, we show the coverage of the mappings for each of the three replicates (top) and the mappings of the respective reads merged across all replicates. Colour indicates the strand orientation of the mapped read pair (blue: forward strand; red: reverse strand). In addition, track “PXO.gff” provides genome-wide predictions of TALE target boxes. Expression of LOC_Os09g29820.1 is likely induced by TalAR3, where the predicted target box obtained rank 41 among all genome-wide predicted target boxes of TalAR3. After *Xoo* PXO83 infection, we observe expression levels increasing over time. The transcription start site appears to be shifted after *Xoo* PXO83 infection relative to the remaining experiments, which can also be perceived from the respective coverage profiles.

### Identification of TALE targets with low PrediTALE ranks

Given their clear expression patterns, specificity for one pathovar and enrichment of predicted TALE targets, Group 3 of *Xoo* PXO83 and Group 1 of *Xoc* BAI35 (Figure 4) could be seen as prototypes of TALE-induced expression. This raises the question if these clusters probably contain further TALE targets that have been neglected when only considering top 50 PrediTALE predictions. Hence, we further use PrediTALE to predict TALE target boxes with a loose p-value threshold (10^−3^) genome-wide for all TALEs present in the TALomes of *Xoo* PXO83 and *Xoc* BAI35, and manually inspect the promoters of the DETs from those clusters for further target boxes.

Indeed, we find several examples for *Xoo* PXO83 and *Xoc* BAI35 that show pathovar-specific up-regulation and/or shifted TSS relative to the other conditions as detailed in the following. For *Xoo* PXO83, LOC_Os09g28440.1 is predicted as a putative target of TalCA1 with prediction rank 77, which shows a shifted TSS after *Xoo* PXO83 infection. OsSWEET14 (LOC_Os11g31190.1) is a well established target of TalAC-class TALEs (16, 47), which contain an aberrant repeat (48, 49). Here, two alternative, overlapping target boxes are possible due to the aberrant repeat. Although both target boxes are predicted with a very low PrediTALE rank, these have been shown to be functional (32, 47, 48), and one reason might be a synergistic effect on DNA binding affinity by these two overlapping target boxes. Notably, we find another example where two overlapping but low-ranking TalAG4 target boxes are located in the promoter of LOC_Os12g04340.1 and likely result in pathovar-specific up-regulation of this transcript. After *Xoc* BAI35 infection, we also find putative additional TALE targets with LOC_Os02g44680.1 (TalBL13, rank 76) and LOC_Os12g42970.1 (TalCB5, rank 214) showing shifted TSSs, and LOC_Os05g39200.1 (TalBD14, rank 138) showing early and pathovar-specific up-regulation.

Taken together, we were able to confirm one well-known TALE target (OsSWEET14) and to identify five novel TALE targets based on their expression profile and shifted transcription start sites.

### Genome-wide identification of TALE-induced differentially expressed regions

Together with PrediTALE, we also published DerTALE as a method for genome-wide TALE target discovery that is not limited to promoter regions and independent of existing gene annotations (11). Here, we present DerTALEv2 as an extended and refined version of DerTALE. In DerTALEv2, putative differentially expressed regions (DERs) are assigned a p-value based on an F-test. In addition, we implemented several heuristic criteria (e.g., distance between target box and start of DER, strand orientation; cf. Methods) to reduce the number of false positive predictions. In this study, we may further filter by the expressional dynamics of DERs over the time course. Specifically, we consider DERs between *Xoo* PXO83 or *Xoc* BAI35 and Mock inoculation, respectively, that are significant (*p<* 0.05) and are present from 24h or 36h until the 72h time point.

Among the putative TALE targets discovered by DerTALEv2 are many TALE targets that have already been found in the promoter-centric analyses. Here, we focus on putative TALE targets that are not well-represented by the MSU7 annotation, including lncRNAs and antisense transcripts (Table 2).

**Table 2.** List of putatively TALE-dependent differentially expressed regions as predicted using DerTALEv2.

| Strain | TALE | Rank | Coordinates | Remarks |
| --- | --- | --- | --- | --- |
| **homology to mRNA** |  |  |  |  |
| *Xoo* PXO83 | TalBA2 | 88 | Chr4:33663291-33664038 | antisense to LOC_Os04g56470.1; match to EU948741.1 (*Zea mays*) |
| *Xoc* BAI35 | TalBD14 | 200 | Chr11:1825439-1826112 | antisense to LOC_Os11g04400.1; match to XM_052291106.1 (*Oryza glaberrima* KIN17-like protein) |
| *Xoc* BAI35 | TalBF17 | 269 | Chr8:2206260-2209729 | match to XM_026027578.1 (*Oryza sativa*), mRNA |
| *Xoc* BAI35 | TalBF17 | 2 | Chr4:26974317-26976201 | match to XR_008016572.1 (*Oryza glaberrima* glycosyltransferase BC10-like) |
| *Xoc* BAI35 | TalCD5 | 76 | Chr3:2459196-2465189 | overlapping LOC_Os03g05049.1; match to XR_003241406.1 (*Oryza sativa* signal peptidase complex catalytic subunit SEC11A-like) |
| **ncRNAs** |  |  |  |  |
| *Xoo* PXO83 | TalAN3 | 13 | Chr11:24504316-24507454 | match to XR_001541283.1 (*Oryza sativa*, LOC107279433), ncRNA |
| *Xoo* PXO83 | TalAO3 | 383 | Chr7:22546154-22547910 | match to XR_001547425.2 (*Oryza sativa*, LOC107281667), ncRNA; previously reported (11) |
| *Xoc* BAI35 | TalBD14 | 201 | Chr12:1758864-1759672 | antisense to LOC_Os12g04180.1; match to XR_006662266.1 (*Aegilops tauschii*, LOC120962805), ncRNA |
| *Xoc* BAI35 | TalBU14 | 845 | Chr1:37104791-37108363 | match to AB427154.1 (*Oryza sativa*, SINE:OsSN1 DNA) |
| **antisense** |  |  |  |  |
| *Xoo* PXO83 | TalAR3 | 1207 | Chr8:9538729-9539728 | antisense to LOC_Os08g15710.1 (log2-fold change 4.9 at 72h) |
| *Xoc* BAI35 | TalBD14 | 169 | Chr5:27423155-27424584 | antisense to LOC_Os05g47840.1 (log2-fold change -0.5 at 72h) |
| *Xoc* BAI35 | TalBF17 | 137 | Chr2:31793228-31793971 | antisense to LOC_Os02g51910.1 (log2-fold change -0.2 at 72h) |
| *Xoc* BAI35 | TalBF17 | 286 | Chr10:1731955-1733975 | antisense to LOC_Os10g03830.1 (log2-fold change -0.2 at 72h) |
| *Xoc* BAI35 | TalBO14 | 1105 | Chr3:32970561-32973139 | antisense to LOC_Os03g57880.2 (log2-fold change 1.8 at 72h) |
| *Xoc* BAI35 | TalBU14 | 13 | Chr4:7205868-7207057 | antisense to LOC_Os04g13060.1 (log2-fold change 2.0 at 72h) |
| *Xoc* BAI35 | TalBU14 | 811 | Chr3:750783-755321 | antisense to LOC_Os03g02240.1 (log2-fold change 0.3 at 72h) |
| *Xoc* BAI35 | TalCJ5 | 1238 | Chr3:1427725-1428741 | antisense to LOC_Os03g03340.1 (log2-fold change 2.5 at 72h); LOC_Os03g03334.1 (log2-fold change 4.9 at 72h) |
| **shifted TSS** |  |  |  |  |
| *Xoo* PXO83 | TalAB5 | 798 | Chr9:17492490-17503103 | LOC_Os09g28810.1; 6,250 bp |
| *Xoo* PXO83 | TalAO3 | 1222 | Chr7:7940492-7944801 | LOC_Os07g13880.1; 2,260 bp |
| *Xoc* BAI35 | TalCF6 | 106 | Chr3:27466967-27472806 | LOC_Os03g48270.1; -390 bp |
| **no match** |  |  |  |  |
| *Xoc* BAI35 | TalAX14 | 687 | Chr1:40340832-40342769 |  |
| *Xoc* BAI35 | TalBO14 | 688 | Chr9:13934613-13935744 |  |
| *Xoc* BAI35 | TalCF6 | 85 | Chr11:27213248-27214914 |  |
| *Xoc* BAI35 | TalCP7 | 154 | Chr5:23981594-23982530 |  |

First, we find several transcripts with BLAST hits to mRNAs from other species or rice mRNAs absent from MSU7. In Supplementary Figure S9 we present an IGV snapshot for one representative example of such cases, where transcription is likely induced by TalBF17 of *Xoc* BAI35.

Second, we find several putatively TALE-induced transcripts with matches to non-coding RNAs, one of which (Chr7:22546154-22547910, predicted target of TalAO3 of *Xoo* PXO83) has been reported previously (11). An IGV snapshot of an additional example (Chr1:37104791-37108363, predicted target of TalBU14 of *Xoc* BAI35) is provided as Supplementary Figure S10. Together, such transcript with matches to known mRNAs or ncRNAs highlight that the current genome annotation of rice is still incomplete.

Third, we frequently observe differentially expressed transcripts overlapping annotated MSU7 transcripts, but that are transcribed in antisense orientation as illustrated in Supplementary Figure S11 for an antisense transcript overlapping LOC_Os05g47840.1 (IPP transferase). These antisense transcripts could have regulatory function, but we do not observe a clear regulatory pattern for the overlapping sense transcripts (Table 2), where some of the sense transcripts are up-regulated and others are (slightly) down-regulated at 72h compared with Mock inoculation.

Fourth, we find additional examples of – in some cases strongly – shifted TSSs, which is the reason why these have been missing in the promoter-centric analysis. In Supplementary Figure S12, we provide an IGV snapshot of the most extreme example (LOC_Os09g28810.1), which is predicted to be induced by TalAB5 of *Xoo* PXO83 and contains an additional upstream region of approx. 6,250 bp compared with the natural transcript present after *Xoc* BAI35 infection or Mock inoculation.

Finally, we find a number of putatively TALE-induced transcripts for which no matching genes (coding or non-coding) exist in NCBI Genbank, and one example of such transcripts is given in Supplementary Figure S13. Whether such targets serve a function during infection or are rather “collateral targets” of TALEs remains unclear without further experiments.

### TALE clusters

TALE genes occur in clusters of up to 5 TALEs (Cluster IX) in *Xoo* PXO83 and up to 6 TALEs in *Xoc* BAI35 (3, 22). Notable differences exist in cluster organization between *X. oryzae* pv. *oryzae* with larger spacers containing inverted repeats and *X. oryzae* pv. *oryzicola* with shorter spacers (50). Here, we can only use the up-regulation of TALE targets as a proxy of the presence of the respective TALE genes. Figure 6 shows the number of predicted target genes of each TALE that are significantly up-regulated from the respective time point until 72h. In general, we find that the number of putative targets that are up-regulated increases over time, which is also reflected on the level of log2-fold changes (Supplementary Figure S14). For *Xoo* PXO83, we find no putative targets for TalAI4 and only one target for TalAI3. Since both are iTALEs/truncTALE (51, 52), these lack the activation domain and, hence, are unable to (directly) activate the expression of target genes. Hence, the lack of targets for TalAI4 is expected and the one prediction for TalAI3 is considered a false-positive. For the remaining TALEs, we observe that some TALEs organized as “singletons” (TalAS3), at the borders of clusters (TalBA2) or contiguously within a cluster (TalAO3/TalAE4/TalAD5) do not have predicted targets at 24h but only 36h after *Xoo* PXO83 infection. However, considering DerTALEv2 predictions S15 individual predictions for some of these TALEs are found at 24h. Considering promoter-based PrediTALE predictions (Figure 6B) and genome-wide DerTALEv2 predictions (Supplementary Figure S15) for *Xoc* BAI35, we do not find targets 24h after infection for TalAV14 (cluster III), TalAT14 (cluster IX), TalAZ11 (cluster VII), TalBI14 (cluster V), and TalCC7. The latter is a pseudo gene and, hence, expected to be dysfunctional in general. We also observe that – especially for *Xoc* BAI35– targets of some TALEs are better reflected by the promoter-centric predictions of PrediTALE, while others are more abundant among the genome-wide DerTALEv2 predictions.

**Fig. 6.**
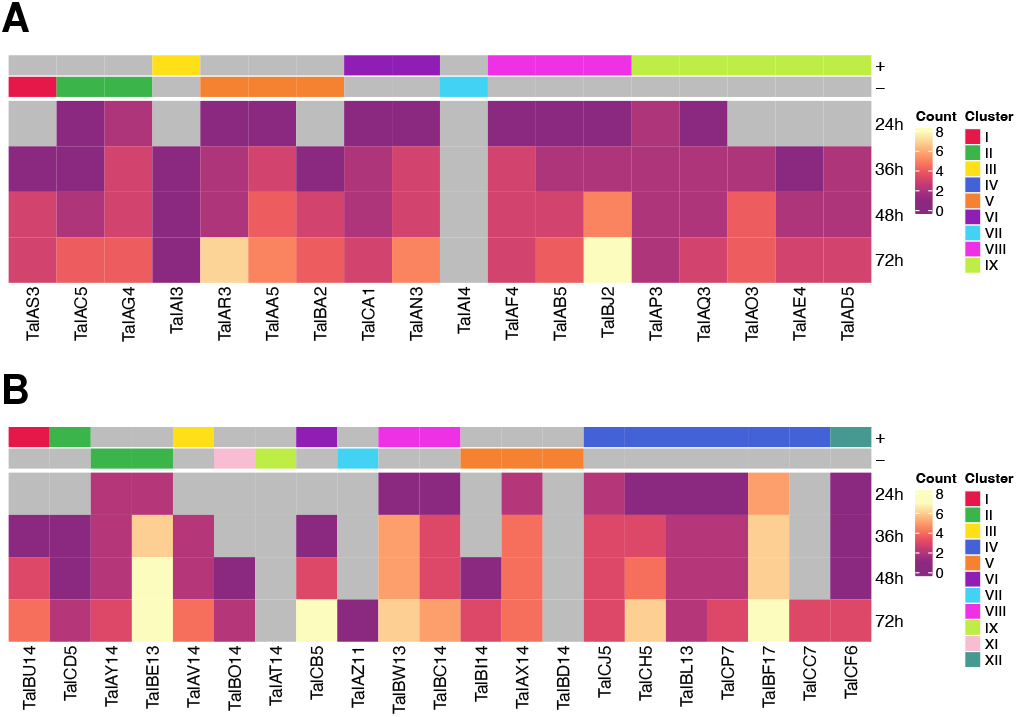
Heatmap visualization of the number of putative target genes per TALE after *Xoo* PXO83 (A) and *Xoc* BAI35 (B) infection. TALEs are ordered by their genomic organization and the respective TALE clusters and strand orientation are indicated by color annotations above the heatmaps. Here, a TALE target is counted if it is consistently up-regulated from the time point indicated at the ordinate until 72h.

### Secondary targets

Time-resolved RNA-seq data after *Xanthomonas* infection provide the unique opportunity to gain further insights into putative secondary TALE targets, i.e. transcripts that are regulated by transcription factors (TFs) that are direct TALE targets. Here, we consider TFs that have a predicted TALE target box in their promoter and that are significantly (adjusted p-value *<* 0.05) up-regulated at least 2-fold (log2-fold change *>* 1) from 24h or 36h until 72h after *Xoo* PXO83 or *Xoc* BAI35 infection, respectively. Identification of putative secondary targets of such TFs may follow different routes, namely (a) based on TF-specific ChIP-seq data where peaks are associated with downstream transcripts, (b) based on known binding motifs and subsequent prediction of binding sites within promoter regions, and (c) based on gene regulatory networks or co-expression networks. In principle, we follow all three routes, but feasibility depends on the availability of the respective data.

### Approaches for identifying secondary targets

For *Xoo* PXO83, we find five TFs (Table 3) fulfilling above criteria: the well-known TALE target OsTFX1 (17), two WRKY TFs, one homeobox and one B-box zinc finger TF. ChIP-seq data is available for OsTFX1 (33), while binding motifs are available for OsWRKY15, OsWRKY24, LOC_Os01g63510.1, and OsTFX1. First, we consider the ChIP-seq peaks for OsTFX1, which have been determined from ChIP-seq data of Flag-fusion protein in an overexpression rice line (33). Specifically, we evaluate the expression of ChIP-associated genes listed in Supplementary Data 2 of (33), ChIP-seq peaks and associated genes called from the same raw data using the unified pipeline of ChIP-Hub (34) and peaks called from the same raw data using our own MACS-based pipeline (cf. Methods). In all three cases, the number of ChIP-associated transcripts that are up-regulated or down-regulated according to our time-resolved RNA-seq data is low. Of the 780 transcripts corresponding to the genes listed in (33), only 27 are up-regulated and only 7 are down-regulated at any time point within 36h to 72h after *Xoo* PXO83 infection. Notably, up-regulation is observed only for one of the peroxidase percursors (LOC_Os03g32050.1) and neither OsNCED3 (LOC_Os03g44380.1) nor OsNCED5 (LOC_Os12g42280.1) where found to be de-regulated in our experiments, although the interaction partner bZIP71 (LOC_Os09g13570.1) of OsTFX1/bZIP73 is strongly expressed (Supplementary Table D). One explanation might be that overexpression and cold treatment that have been the focus of (33), do not reflect the function of OsTFX1 in the *Xanthomonas* infection. ChIP-associated transcripts from ChIP-Hub (19 up-regulated and 6 down-regulated of 522 ChIP-associated transcripts in total) and from our own MACS-based pipeline (12 up-regulated and 6 down-regulated of 707 ChIP-associated transcripts in total) reflect the results based on the gene list from the original publication.

**Table 3.** Transcription factor TALE targets up-regulated after *Xoo* PXO83 and *Xoc* BAI35 infection. Columns “ChIP-seq” and “motif” indicate the availability of the respective resources for a TF. Log2-fold changes (columns with prefix “lfc”) are provided for the infection with the respective strain relative to Mock inoculation.

| Strain | TALE | Transcript | Annotation | ChIP-seq | motif | lfc 24h | lfc 36h | lfc 48h | lfc 72h |
| --- | --- | --- | --- | --- | --- | --- | --- | --- | --- |
| *Xoo* PXO83 | TalAA5 | LOC_Os01g46800.1 | OsWRKY15 | no | yes | 2.4 | 1.5 | 2.5 | 2.9 |
| *Xoo* PXO83 | TalAS3 | LOC_Os01g61080.1 | OsWRKY24 | no | yes | 1.9 | 2.5 | 2.7 | 3.2 |
| *Xoo* PXO83 | TalCA1 | LOC_Os01g63510.1 | homeobox domain | no | yes | 6.7 | 7.6 | 9.2 | 10.2 |
| *Xoo* PXO83 | TalBJ2 | LOC_Os04g45690.1 | B-box zinc finger | no | no | 0.2 | 1.0 | 1.9 | 1.9 |
| *Xoo* PXO83 | TalAR3 | LOC_Os09g29820.1 | OsTFX1 | yes | yes | 4.0 | 6.9 | 4.6 | 4.6 |
| *Xoc* BAI35 | TalCD5 | LOC_Os01g39330.1 | OsBHLH024 | no | no | 0.6 | 2.4 | 1.2 | 1.6 |
| *Xoc* BAI35 | TalBF17 | LOC_Os08g37400.1 | OsZHD2 | no | no | 1.3 | 3.2 | 2.2 | 2.5 |

For four of the TFs that are putative targets of *Xoo* PXO83 TALEs, we further consider motifs from CIS-BP (38) and PlantTFDB (53) as listed in Supplementary Table C. Here, we predict binding sites based on the binding motifs using Fimo (40) with a significance threshold of 10^−5^. The number of up-regulated or down-regulated transcripts among those with a predicted binding site of the respective TF according to the Fimo prediction is slightly larger. For OsWRKY15, we find 46 up-regulated and 4 down-regulated transcripts 72h after *Xoo* PXO83 infection of 633 targets predicted by Fimo in total; for OsWRKY24, we find 46 up-regulated and 2 down-regulated transcripts of 713 in total; for LOC_Os01g63510.1, we find 31 up-regulated and 12 down-regulated transcripts of 686 in total; and for OsTFX1, we find 43 up-regulated and 18 down-regulated transcripts of 1854 in total.

Finally, we consider co-expression networks generated using the arboreto re-implementation (43) of the GENIE3 method (42) based on 785 publicly available RNA-seq data sets (cf. Methods). The main advantage of this approach is that it can be conducted for all TFs listed in Table 3, although co-expression is only a rough proxy for (direct) regulatory interactions. Here, we consider the top 500 co-expressed transcripts per TF to yield numbers that are comparable to previous ChIP-based and motif-based analyses. Of the top 500 transcripts co-expressed with OsWRKY15, 80 are up-regulated and 3 are down-regulated at any time point within 36h to 72h after *Xoo* PXO83 infection; for OsWRKY24, we find 211 up-regulated and 1 down-regulated transcript(s); for LOC_Os01g63510.1, we find 3 up-regulated and 1 down-egulated transcript(s); for LOC_Os04g45690.1, we find 38 up-regulated and 7 down-regulated transcripts; and for OsTFX1, we find 14 up-regulated and 1 down-regulated transcript(s).

The concordance of the different approaches is also low. For OsTFX1, we find one overlapping transcript between the ChIP-associated and the motif-based transcripts, no overlap between the ChIP-associated and the network-based transcripts, and one overlapping transcripts between the motif-based and the network-based transcripts. For OsWRKY15, we find 3 overlapping transcripts based on motifs and networks; for OsWRKY24 8 overlapping transcripts, and for LOC_Os01g63510.1 no overlapping transcripts.

With the network-based approach, we may further determine putative targets of the TFs that are putative direct targets of *Xoc* BAI35 TALEs. We find 238 up-regulated and 1 down-regulated transcripts among those co-expressed with Os-bHLH024, and 13 up-regulated and 11 down-regulated transcripts for OsZHD2. In the following, we refer to TFs that are (direct) TALE targets as “TALE-induced TFs”, whereas we refer to (putative) regulatory target transcripts of TFs as “TF targets”.

Being aware of its limitations but due to lack of alternatives, we consider the putative TF targets identified by the network-based approach in the following, since it is also applicable to the TFs induced after *Xoc* BAI35 infection and yields the largest number of putative TF targets in total.

### Clustering direct and secondary targets

In Figure 7 A and B, we present cluster plots of the expression profiles of putative TALE and TF targets after *Xoo* PXO83 and *Xoc* BAI35 infection, respectively. Here, we consider all transcripts that are up-regulated at any of the time points considered, since the dynamics of TF-induced transcription may be variable. For *Xoo* PXO83, we find one cluster (Group 4) that contains equal numbers of up-regulated transcripts that are predicted to be induced by TALEs or TFs. In both cases, up-regulation is specific for the *Xoo* PXO83 infection and expression levels after *Xoc* BAI35 infection is similar to Mock inoculation. By contrast, the expression in two large clusters (Group 1 and Group 3) that contain a larger number of TF-induced than TALE-induced transcripts, up-regulation can be observed after *Xoc* BAI35 infection as well. In one smaller cluster (Group 2), induction is shifted towards later time points.

**Fig. 7.**
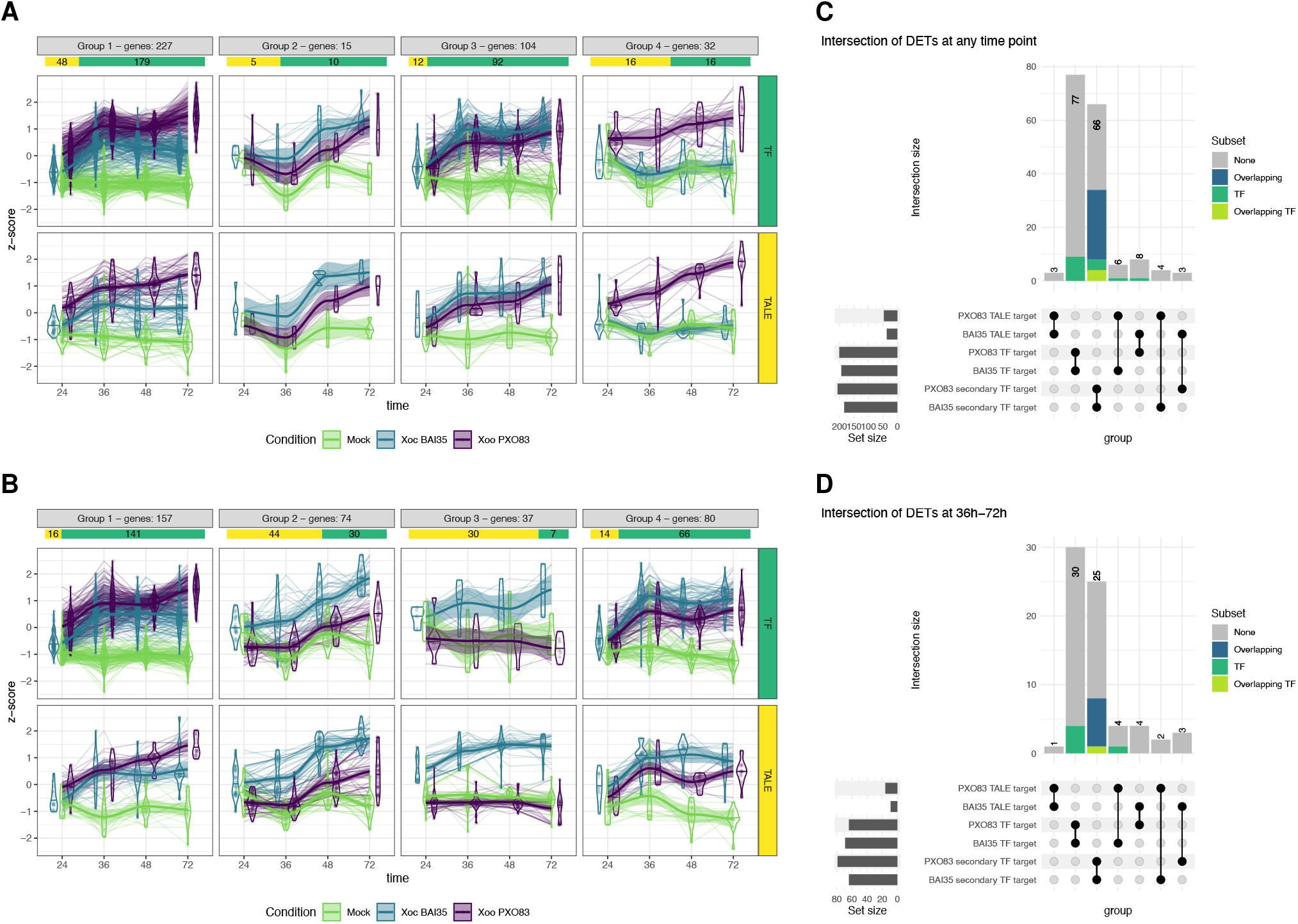
Clustered expression profiles of putative TALE targets and putative TF targets after *Xoo* PXO83 (A) and *Xoc* BAI35 (B) infection. Proportions of TALE targets and TF targets within each cluster are visualized below the respective cluster title bar. For each cluster, we separately plot expression profiles of putative TALE targets (bottom) and putative TF targets (top). (C) Upset plot of direct TALE targets, targets of TALE-induced TFs, and secondary TF targets, i.e. targets of TFs that are activated by TALE-induced TFs. Here, we consider all transcripts that are up-regulated at at least one time point. Putative TF targets have been determined based on co-expression networks. Only a subset of all possible intersections is shown for clarity. Within the bar for each intersection, we highlight transcripts that are TFs. We further indicate (“Overlapping”) if a transcript has been activated by one of the TFs in the intersection between “PXO83 TF target” and “BAI35 TF target”. (D) Same as (C) but for transcripts that are consistently up-regulated from 36h to 72h after the infection.

For *Xoc* BAI35 (Figure 7B), we find one cluster (Group 3) that is dominated by putative TALE targets and, again, shows a transcriptional response specific for *Xoc* BAI35 infection. Group 2 also contains a larger number of TALE-induced than TF-induced transcripts and shows similar tendencies but with a later response. The largest cluster (Group 1), in turn, is dominated by TF-induced transcripts and, again, shows similar response after *Xoo* PXO83 and *Xoc* BAI35 infection, while the remaining cluster (Group 4) contains transcripts with a stronger response to *Xoc* BAI35 infection, but up-regulation also present after *Xoo* PXO83 infection.

The results of the clustering indicate that TALE-dependent induction of expression may be rather specific to one of the pathovars, whereas TF-dependent induction has the tendency to target transcripts that are commonly up-regulated after *Xoo* PXO83 and *Xoc* BAI35 infection.

### GO terms enriched for secondary targets

We further determine GO terms that are enriched among the putative TF-induced transcript after *Xoo* PXO83 and *Xoc* BAI35 infection, respectively, as shown in Supplementary Figure S16. For the “molecular function” ontology, we observe – besides general terms – “kinase activity”, “protein binding” and “transcription factor activity” enriched at several time points after *Xoo* PXO83 and *Xoc* BAI35 infection. These may be associated with signal transduction and stress response and, indeed, we find those terms enriched for the “biological process” ontology, but also “response to endogenous stimulus” and “response to biotic stimulus”. For the “cellular component” ontology, the term “plasma membrane” is significantly enriched for *Xoo* PXO83 but not for *Xoc* BAI35.

### Downstream targets of TFs

Since the GO term “transcription factor activity” has been found to be enriched among the putative TF targets it might be that further TFs are induced indirectly via TALE-induced TFs. Hence, we screen those TF targets for further TF transcripts. We find 34 further TFs for *Xoo* PXO83 and 23 further TFs for *Xoc* BAI35, 9 of which are shared between both pathovars. If we limit the set of transcripts considered to those that are already up-regulated at 36h or 48h, we still find 30 further TFs for *Xoo* PXO83 and 19 further TFs for *Xoc* BAI35, 8 of which are shared. Again, we query the co-expression network for putative targets of these TFs, but limit the number of candidates per TF to 100 to keep the numbers comparable to the previously determined TF targets. Due to the overlap of indirect TF targets (intersection between “PXO83 TF target” and “BAI35 TF target” in Figure 7C) between the pathovars, such secondary TF targets will overlap to a substantial extent. In Figure 7C, we visualize the overlap between direct TALE targets, direct TF targets, and secondary TF targets between the two pathovars considering transcripts that are up-regulated at any of the time points (cf. panel A and B). We find only 3 overlapping transcripts between the *Xoo* PXO83 and *Xoc* BAI35 TALE targets, namely the previously mentioned expressed protein (LOC_Os01g40290.1), LOC_Os01g33090.1 and LOC_Os02g48570.1. In the intersection between *Xoo* PXO83 and *Xoc* BAI35 TF targets, we find 77 transcripts including the 9 shared TFs, which indicates that the pathogen-specific direct up-regulation by TALEs becomes more common between *Xoo* PXO83 and *Xoc* BAI35 on the level of secondary TALE targets. For secondary TF targets, we observe 66 transcripts shared between *Xoo* PXO83 and *Xoc* BAI35, 30 of which are putatively induced by the TFs shared between both pathovars. In addition, some transcripts are putatively up-regulated directly by a TALE in one pathovar, but indirectly via a TF in the other pathovar. Notably, this includes OsWRKY24, which is predicted as a direct target of TalAS3 is *Xoo* PXO83 but a putative indirect target via OsbHLH024 in *Xoc* BAI35 and – vice versa – OsbHLH024 is predicted as a direct target of TalCD5 in *Xoc* BAI35 but a putative indirect target via OsWRKY24 in *Xoo* PXO83. However, the basis of these analyses is a co-expression network and, hence, OsWRKY24 and OsbHLH024 are co-expressed in general. Limiting the transcripts considered to those up-regulated from 36h to 72h (Figure 7D) lowers the numbers in the intersections, but the general picture remains similar.

In summary, our findings support a model where the same downstream, secondary targets are induced via different direct TALE targets in different pathovars or strains. By contrast, we find limited evidence that targets may be induced directly via TALEs in one strain and indirectly as secondary targets in another strain, or that both activation pathways are used interchangeably.

## Conclusions

The time-resolved expression data after *Xoo* PXO83 and *Xoc* BAI35 infection provide new insights into the dynamics of TALE-dependent activation of target transcripts. Based on the findings presented in this manuscript, we formulate four criteria that may help to identify true TALE targets:

1. Consistent up-regulation from early (24h or 36h) until late (72h) time points.
2. Apparently accumulating expression levels over time that are observed only after the infection but not in Mock inoculation. These may be either an effect of continuous activation by TALEs or by increasing numbers of infected plant cells.
3. Specific activation only in strains harbouring the respective TALE. We find little evidence of alternative activation pathways (directly via TALEs or as secondary targets via TFs that are TALE targets themselves). Exceptions from this rule are common targets that are induced by different TALEs in *Xoo* and *Xoc*, respectively (cf. LOC_Os01g40290.1).
4. Shifted transcription start sites relative to the location of the TALE target box. If those are observed consistently across time points, they are a strong indication of TALE-dependent activation. However, depending on the distance of the TALE target box to the natural TSS, TALE-induced transcription may also start from the natural TSS.

Except for the first criterion, these criteria do not apply to all TALE targets. However, the larger the number of criteria fulfilled, the stronger do we consider the evidence that the expression of a transcript up-regulated after the infection is indeed a TALE target.

We further identified TALE targets based on genome-wide target box predictions independent of existing gene annotations using DerTALEv2. Our findings suggest that currently incomplete gene annotations of the rice genome are a limiting factor of promoter-centric approaches. In addition, we find a considerable number of TALE-induced non-coding RNAs and antisense transcripts that may have regulatory function. We further conclude that the identification of secondary TALE targets, i.e. target genes that are regulated by TFs that are direct TALE targets, remains complex even based on time-resolved expression data. To reliably identify such secondary targets, further experiments about TF binding (ChIP-seq, DAP-seq) are required and – at least for OsTFX1 considered in this study – ChIP-seq data collected under different experimental conditions are insufficient for this purpose. Taken together, our study based on co-expression networks gives first indications that secondary targets contribute to converging sets of up-regulated transcripts after *Xoo* PXO83 and *Xoc* BAI35 infection.

## DATA AVAILABILITY

Raw sequencing data of all RNA-seq libraries are available from the European Nucleotide Archive (https://www.ebi.ac.uk/ena/) under Accession PRJEB76065. The binary version of DerTALEv2 is available from https://www.jstacs.de/index.php/DerTALEv2. Source code is available from the Jstacs GitHub-Repository at https://github.com/Jstacs/Jstacs; code specific to Der-TALEv2 is located at https://github.com/Jstacs/Jstacs/tree/master/projects/tals/rnaseq.

## FUNDING

This work was supported by grants from the Deutsche Forschungsgemeinschaft (http://www.dfg.de) (BO 1496/8-2 to JB and GR 4587/1-2 to JG). The funders had no role in study design, data collection and analysis, decision to publish, or preparation of the manuscript.

This preprint is formatted using a LATEX class by Ricardo Henriques.

## REFERENCES

1. Wende Liu, Jinling Liu, Lindsay Triplett, Jan E. Leach, and Guo-Liang Wang. Novel insights into rice innate immunity against bacterial and fungal pathogens. Annu Rev Phytopathol, 52 (1):213–241, 2014.

2. Jens Boch and Ulla Bonas. Xanthomonas AvrBs3 family-type III effectors: discovery and function. Annu Rev Phytopathol, 48(1):419–436, 2010. doi: 10.1146/annurev-phyto-080508-081936.

3. Jan Grau, Maik Reschke, Annett Erkes, Jana Streubel, Richard D. Morgan, Geoffrey G. Wilson, Ralf Koebnik, and Jens Boch. AnnoTALE: bioinformatics tools for identification, annotation, and nomenclature of tales from xanthomonas genomic sequences. Scientific Reports, 6:21077 EP –, 02 2016.

4. Jens Boch, Heidi Scholze, Sebastian Schornack, Angelika Landgraf, Simone Hahn, Sabine Kay, Thomas Lahaye, Anja Nickstadt, and Ulla Bonas. Breaking the code of DNA binding specificity of TAL-type III effectors. Science, 326(5959):1509–1512, 2009. doi: 10.1126/science.1178811.

5. Matthew J. Moscou and Adam J. Bogdanove. A simple cipher governs DNA recognition by TAL effectors. Science, 326(5959):1501–1501, 2009. doi: 10.1126/science.1178817.

6. Amanda Nga-Sze Mak, Philip Bradley, Raul A. Cernadas, Adam J. Bogdanove, and Barry L. Stoddard. The crystal structure of TAL effector PthXo1 bound to its DNA target. Science, 335(6069):716–9, 2012.

7. Junjiao Yang, Yuan Zhang, Pengfei Yuan, Yuexin Zhou, Changzu Cai, Qingpeng Ren, Dingqiao Wen, Coco Chu, Hai Qi, and Wensheng Wei. Complete decoding of TAL effectors for DNA recognition. Cell Res, 24(5):628–631, 2014.

8. Jan Grau, Annett Wolf, Maik Reschke, Ulla Bonas, Stefan Posch, and Jens Boch. Computational predictions provide insights into the biology of TAL effector target sites. PLoS Comput Biol, 9(3):e1002962, 2013. doi: 10.1371/journal.pcbi.1002962.

9. Erin L. Doyle, Nicholas J. Booher, Daniel S. Standage, Daniel F. Voytas, Volker P. Brendel, John K. VanDyk, and Adam J. Bogdanove. TAL effector-nucleotide targeter (TALE-NT) 2.0: tools for TAL effector design and target prediction. Nucleic Acids Res, 40(W1):W117–W122, 2012.

10. Alvaro L. Pérez-Quintero, Luis M. Rodriguez-R, Alexis Dereeper, Camilo López, Ralf Koebnik, Boris Szurek, and Sebastien Cunnac. An improved method for TAL effectors DNA-binding sites prediction reveals functional convergence in TAL repertoires of Xanthomonas oryzae strains. PLoS ONE, 8(7):e68464.EP, 2013.

11. Annett Erkes, Stefanie Mücke, Maik Reschke, Jens Boch, and Jan Grau. PrediTALE: A novel model learned from quantitative data allows for new perspectives on TALE targeting. PLoS Comput Biol, 15(7):1–31, 2019. doi: 10.1371/journal.pcbi.1007206.

12. Dong Deng, Ping Yin, Chuangye Yan, Xiaojing Pan, Xinqi Gong, Shiqian Qi, Tian Xie, Magdy Mahfouz, Jian-Kang Zhu, Nieng Yan, and Yigong Shi. Recognition of methylated DNA by TAL effectors. Cell Res, 22(10):1502–1504, 2012.

13. Annett Erkes, Stefanie Mücke, Maik Reschke, Jens Boch, and Jan Grau. Epigenetic features improve tale target prediction. BMC Genomics, 22(1):914, 2021. doi: 10.1186/s12864-021-08210-z.

14. Katherine Wilkins, Nicholas Booher, Li Wang, and Adam Bogdanove. TAL effectors and activation of predicted host targets distinguish asian from african strains of the rice pathogen Xanthomonas oryzae pv. oryzicola while strict conservation suggests universal importance of five TAL effectors. Front Plant Sci, 6:536, 2015.

15. Zhou-Xiang Liao, Zhe Ni, Xin-Li Wei, Long Chen, Jian-Yuan Li, Yan-Hua Yu, Wei Jiang, BoLe Jiang, Yong-Qiang He, and Sheng Huang. Dual RNA-seq of Xanthomonas oryzae pv. oryzicola infecting rice reveals novel insights into bacterial-plant interaction. PLOS ONE, 14 (4):1–16, 04 2019. doi: 10.1371/journal.pone.0215039.

16. Stefanie Mücke, Maik Reschke, Annett Erkes, Claudia-Alice Schwietzer, Sebastian Becker, Jana Streubel, Richard D. Morgan, Geoffrey G. Wilson, Jan Grau, and Jens Boch. Transcriptional reprogramming of rice cells by Xanthomonas oryzae TALEs. Frontiers in Plant Science, 10, 2019. ISSN 1664-462X. doi: 10.3389/fpls.2019.00162.

17. Akiko Sugio, Bing Yang, Tong Zhu, and Frank F. White. Two type iii effector genes of Xanthomonas oryzae pv. oryzae control the induction of the host genes OsTFIIA”1 and OsTFX1 during bacterial blight of rice. Proceedings of the National Academy of Sciences, 104 (25):10720–10725, 2007. doi: 10.1073/pnas.0701742104.

18. Tuan T. Tran, Alvaro L. Pérez-Quintero, Issa Wonni, Sara C. D. Carpenter, Yanhua Yu, Li Wang, Jan E. Leach, Valérie Verdier, Sébastien Cunnac, Adam J. Bogdanove, Ralf Koebnik, Mathilde Hutin, and Boris Szurek. Functional analysis of African Xanthomonas oryzae pv. oryzae TALomes reveals a new susceptibility gene in bacterial leaf blight of rice. PLOS Pathogens, 14(6):1–25, 06 2018. doi: 10.1371/journal.ppat.1007092.

19. Rezwan Tariq, Zhiyuan Ji, Chunlian Wang, Yongchao Tang, Lifang Zou, Hongda Sun, Gongyou Chen, and Kaijun Zhao. RNA-seq analysis of gene expression changes triggered by Xanthomonas oryzae pv. oryzae in a susceptible rice genotype. Rice, 12(1):44, 2019. doi: 10.1186/s12284-019-0301-2.

20. Chunlian Wang, Rezwan Tariq, Zhiyuan Ji, Zheng Wei, Kaili Zheng, Rukmini Mishra, and Kaijun Zhao. Transcriptome analysis of a rice cultivar reveals the differentially expressed genes in response to wild and mutant strains of xanthomonas oryzae pv. oryzae. Scientific Reports, 9(1):3757, 2019. doi: 10.1038/s41598-019-39928-2.

21. Rezwan Tariq, Chunlian Wang, Tengfei Qin, Feifei Xu, Yongchao Tang, Ying Gao, Zhiyuan Ji, and Kaijun Zhao. Comparative transcriptome profiling of rice near-isogenic line carrying xa23 under infection of xanthomonas oryzae pv. oryzae. International Journal of Molecular Sciences, 19(3), 2018. ISSN 1422-0067. doi: 10.3390/ijms19030717.

22. Annett Erkes, RenéP. Grove, Milena Žarković, Sebastian Krautwurst, Ralf Koebnik, Richard D. Morgan, Geoffrey G. Wilson, Martin Hölzer, Manja Marz, Jens Boch, and Jan Grau. Assembling highly repetitive Xanthomonas TALomes using Oxford Nanopore sequencing. BMC Genomics, 24(1):151, 2023. doi: 10.1186/s12864-023-09228-1.

23. David O. Niño-Liu, Pamela C. Ronald, and Adam J. Bogdanove. Xanthomonas oryzae pathovars: model pathogens of a model crop. Molecular Plant Pathology, 7(5):303–324, 2006. doi: 10.1111/j.1364-3703.2006.00344.x.

24. Frank F. White and Bing Yang. Host and Pathogen Factors Controlling the Rice-Xanthomonas oryzae Interaction. Plant Physiology, 150(4):1677–1686, 05 2009. ISSN 0032-0889. doi: 10.1104/pp.109.139360.

25. Alexander Dobin, Carrie A. Davis, Felix Schlesinger, Jorg Drenkow, Chris Zaleski, Sonali Jha, Philippe Batut, Mark Chaisson, and Thomas R. Gingeras. STAR: ultrafast universal RNA-seq aligner. Bioinformatics, 29(1):15–21, 10 2012. ISSN 1367-4803. doi: 10.1093/bioinformatics/bts635.

26. R Core Team. R: A Language and Environment for Statistical Computing. R Foundation for Statistical Computing, Vienna, Austria, 2024.

27. Michael I. Love, Wolfgang Huber, and Simon Anders. Moderated estimation of fold change and dispersion for RNA-seq data with DESeq2. Genome Biology, 15:550, 2014. doi: 10.1186/s13059-014-0550-8.

28. Jens Keilwagen, Michael Wenk, Jessica L. Erickson, Martin H. Schattat, Jan Grau, and Frank Hartung. Using intron position conservation for homology-based gene prediction. Nucleic Acids Research, 44(9):e89–e89, 02 2016. ISSN 0305-1048. doi: 10.1093/nar/gkw092.

29. Jens Keilwagen, Frank Hartung, Michael Paulini, Sven O. Twardziok, and Jan Grau. Combining RNA-seq data and homology-based gene prediction for plants, animals and fungi. BMC Bioinformatics, 19(1):189, 2018. doi: 10.1186/s12859-018-2203-5.

30. Helga Thorvaldsdóttir, James T. Robinson, and Jill P. Mesirov. Integrative Genomics Viewer (IGV): high-performance genomics data visualization and exploration. Briefings in Bioinformatics, 14(2):178–192, 04 2012. ISSN 1467-5463. doi: 10.1093/bib/bbs017.

31. James T Robinson, Helga Thorvaldsdóttir, Wendy Winckler, Mitchell Guttman, Eric S Lander, Gad Getz, and Jill P Mesirov. Integrative genomics viewer. Nature Biotechnology, 29 (1):24–26, 2011. doi: 10.1038/nbt.1754.

32. Jana Streubel, Heidi Baum, Jan Grau, Johannes Stuttman, and Jens Boch. Dissection of TALE-dependent gene activation reveals that they induce transcription cooperatively and in both orientations. PLOS ONE, 12(3):1–24, 03 2017. doi: 10.1371/journal.pone.0173580.

33. Citao Liu, Shujun Ou, Bigang Mao, Jiuyou Tang, Wei Wang, Hongru Wang, Shouyun Cao, Michael R. Schläppi, Bingran Zhao, Guoying Xiao, Xiping Wang, and Chengcai Chu. Early selection of bzip73 facilitated adaptation of japonica rice to cold climates. Nature Communications, 9(1):3302, 2018. doi: 10.1038/s41467-018-05753-w.

34. Liang-Yu Fu, Tao Zhu, Xinkai Zhou, Ranran Yu, Zhaohui He, Peijing Zhang, Zhigui Wu, Ming Chen, Kerstin Kaufmann, and Dijun Chen. ChIP-Hub provides an integrative platform for exploring plant regulome. Nature Communications, 13(1):3413, 2022. doi: 10.1038/s41467-022-30770-1.

35. Ben Langmead and Steven L. Salzberg. Fast gapped-read alignment with Bowtie 2. Nat Methods, 9(4):357–359, 2012.

36. Yong Zhang, Tao Liu, Clifford A. Meyer, Jérôme Eeckhoute, David S. Johnson, Bradley E. Bernstein, Chad Nusbaum, Richard M. Myers, Myles Brown, Wei Li, and X. Shirley Liu. Model-based analysis of ChIP-seq (MACS). Genome Biology, 9(9):R137, 2008. doi: 10.1186/gb-2008-9-9-r137.

37. Aaron R. Quinlan and Ira M. Hall. BEDTools: a flexible suite of utilities for comparing genomic features. Bioinformatics, 26(6):841–842, 01 2010. ISSN 1367-4803. doi: 10.1093/bioinformatics/btq033.

38. Matthew T. Weirauch, Ally Yang, Mihai Albu, Atina G. Cote, Alejandro Montenegro-Montero, Philipp Drewe, Hamed S. Najafabadi, Samuel A. Lambert, Ishminder Mann, Kate Cook, Hong Zheng, Alejandra Goity, Harm van Bakel, Jean-Claude Lozano, Mary Galli, Mathew G. Lewsey, Eryong Huang, Tuhin Mukherjee, Xiaoting Chen, John S. Reece-Hoyes, Sridhar Govindarajan, Gad Shaulsky, Albertha J. M. Walhout, François-Yves Bouget, Gunnar Ratsch, Luis F. Larrondo, Joseph R. Ecker, and Timothy R. Hughes. Determination and inference of eukaryotic transcription factor sequence specificity. Cell, 158(6):1431–1443, 2024/05/14 2014. doi: 10.1016/j.cell.2014.08.009.

39. Jan Grau, Stefan Posch, Ivo Grosse, and Jens Keilwagen. A general approach for discriminative de novo motif discovery from high-throughput data. Nucleic Acids Research, 41(21): e197, 2013. doi: 10.1093/nar/gkt831.

40. Charles E. Grant, Timothy L. Bailey, and William Stafford Noble. FIMO: scanning for occurrences of a given motif. Bioinformatics, 27(7):1017–1018, 02 2011. ISSN 1367-4803. doi: 10.1093/bioinformatics/btr064.

41. Nicolas L Bray, Harold Pimentel, Páll Melsted, and Lior Pachter. Near-optimal probabilistic rna-seq quantification. Nature Biotechnology, 34(5):525–527, 2016. doi: 10.1038/nbt.3519.

42. Vân Anh Huynh-Thu, Alexandre Irrthum, Louis Wehenkel, and Pierre Geurts. Inferring regulatory networks from expression data using tree-based methods. PLOS ONE, 5(9): 1–10, 09 2010. doi: 10.1371/journal.pone.0012776.

43. Sara Aibar, Carmen Bravo González-Blas, Thomas Moerman, Vân Anh Huynh-Thu, Hana Imrichova, Gert Hulselmans, Florian Rambow, Jean-Christophe Marine, Pierre Geurts, Jan Aerts, Joost van den Oord, Zeynep Kalender Atak, Jasper Wouters, and Stein Aerts. Scenic: single-cell regulatory network inference and clustering. Nature Methods, 14(11):1083–1086, 2017. doi: 10.1038/nmeth.4463.

44. Yang Liao, Gordon K. Smyth, and Wei Shi. featureCounts: an efficient general purpose program for assigning sequence reads to genomic features. Bioinformatics, 30(7):923–930, 11 2013. ISSN 1367-4803. doi: 10.1093/bioinformatics/btt656.

45. Lorena Pantano. DEGreport: Report of DEG analysis, 2023. R package version 1.36.0.

46. Patrick Römer, Sabine Recht, Tina Strauß, Janett Elsaesser, Sebastian Schornack, Jens Boch, Shiping Wang, and Thomas Lahaye. Promoter elements of rice susceptibility genes are bound and activated by specific TAL effectors from the bacterial blight pathogen, Xanthomonas oryzae pv. oryzae. New Phytologist, 187(4):1048–1057, 2010. doi: 10.1111/j.1469-8137.2010.03217.x.

47. Ginny Antony, Junhui Zhou, Sheng Huang, Ting Li, Bo Liu, Frank White, and Bing Yang. Rice xa13 Recessive Resistance to Bacterial Blight Is Defeated by Induction of the Disease Susceptibility Gene Os-11N3. The Plant Cell, 22(11):3864–3876, 11 2010. ISSN 1040-4651. doi: 10.1105/tpc.110.078964.

48. Annekatrin Richter, Jana Streubel, Christina Blücher, Boris Szurek, Maik Reschke, Jan Grau, and Jens Boch. A TAL effector repeat architecture for frameshift binding. Nat Commun, 5:3447, 2014. doi: 10.1038/ncomms4447.

49. Sebastian Becker, Stefanie Mücke, Jan Grau, and Jens Boch. Flexible TALEs for an expanded use in gene activation, virulence and scaffold engineering. Nucleic Acids Research, 50(4):2387–2400, 02 2022. ISSN 0305-1048. doi: 10.1093/nar/gkac098.

50. Annett Erkes, Maik Reschke, Jens Boch, and Jan Grau. Evolution of transcription activator-like effectors in Xanthomonas oryzae. Genome Biology and Evolution, 9(6):1599–1615, 2017. doi: 10.1093/gbe/evx108.

51. Zhiyuan Ji, Chonghui Ji, Bo Liu, Lifang Zou, Gongyou Chen, and Bing Yang. Interfering TAL effectors of Xanthomonas oryzae neutralize R-gene-mediated plant disease resistance. Nature Communications, 7(1):13435, 2016. doi: 10.1038/ncomms13435.

52. Andrew C. Read, Fabio C. Rinaldi, Mathilde Hutin, Yong-Qiang He, Lindsay R. Triplett, and Adam J. Bogdanove. Suppression of Xo1-mediated disease resistance in rice by a truncated, non-DNA-binding TAL effector of Xanthomonas oryzae. Frontiers in Plant Science, 7, 2016. ISSN 1664-462X. doi: 10.3389/fpls.2016.01516.

53. Feng Tian, De-Chang Yang, Yu-Qi Meng, Jinpu Jin, and Ge Gao. PlantRegMap: charting functional regulatory maps in plants. Nucleic Acids Research, 48(D1):D1104–D1113, 11 2019. ISSN 0305-1048. doi: 10.1093/nar/gkz1020.

