## Supplementary Table A and C, captions of Supplementary Tables B, D-E and Supplementary Data S1 and S2, Supplementary Figures S1 - S16 for "Time-resolved RNA-seq data reveal dynamic expressional behaviour of TALE target genes"

### SUPPLEMENTARY MATERIAL

**Supplementary Table A.** Read-level statistics of the RNA-seq samples after *Xoo* PXO83 and *Xoc* BAI35 infection and mock inoculation.

| ENA Run Accession | Condition | Time | Reads | Uniquely mapped |
|---|---|---|---|---|
| ERR13160312 | Mock | 24h | 49,105,703 | 85.97% |
| ERR13160345 | Mock | 24h | 38,566,914 | 89.32% |
| ERR13160360 | Mock | 24h | 33,359,014 | 87.85% |
| ERR13160320 | Mock | 36h | 36,705,685 | 62.32% |
| ERR13160369 | Mock | 36h | 34,614,107 | 87.14% |
| ERR13160389 | Mock | 36h | 39,544,023 | 85.79% |
| ERR13160329 | Mock | 48h | 30,930,068 | 83.66% |
| ERR13160352 | Mock | 48h | 28,475,897 | 83.45% |
| ERR13160398 | Mock | 48h | 33,222,387 | 85.68% |
| ERR13160337 | Mock | 72h | 34,726,524 | 86.60% |
| ERR13160377 | Mock | 72h | 34,032,619 | 86.22% |
| ERR13160407 | Mock | 72h | 28,023,902 | 84.33% |
| ERR13160415 | PXO83 | 24h | 32,543,942 | 82.06% |
| ERR13160450 | PXO83 | 24h | 28,979,728 | 82.77% |
| ERR13160497 | PXO83 | 24h | 34,217,903 | 85.34% |
| ERR13160423 | PXO83 | 36h | 30,916,857 | 81.72% |
| ERR13160459 | PXO83 | 36h | 33,282,819 | 82.23% |
| ERR13160506 | PXO83 | 36h | 29,953,231 | 79.35% |
| ERR13160431 | PXO83 | 48h | 28,875,297 | 78.87% |
| ERR13160469 | PXO83 | 48h | 33,739,534 | 81.37% |
| ERR13160516 | PXO83 | 48h | 35,279,177 | 84.55% |
| ERR13160441 | PXO83 | 72h | 35,643,199 | 83.21% |
| ERR13160479 | PXO83 | 72h | 38,382,721 | 83.59% |
| ERR13160488 | PXO83 | 72h | 32,688,821 | 79.31% |
| ERR13160213 | BAI35 | 24h | 31,028,067 | 78.38% |
| ERR13160242 | BAI35 | 24h | 29,048,715 | 78.86% |
| ERR13160276 | BAI35 | 24h | 41,619,784 | 82.69% |
| ERR13160221 | BAI35 | 36h | 38,672,018 | 81.42% |
| ERR13160250 | BAI35 | 36h | 30,744,375 | 71.06% |
| ERR13160286 | BAI35 | 36h | 42,777,459 | 79.11% |
| ERR13160228 | BAI35 | 48h | 32,064,700 | 77.93% |
| ERR13160257 | BAI35 | 48h | 27,996,254 | 77.92% |
| ERR13160294 | BAI35 | 48h | 28,518,442 | 82.03% |
| ERR13160235 | BAI35 | 72h | 33,275,448 | 81.66% |
| ERR13160266 | BAI35 | 72h | 36,090,091 | 83.56% |
| ERR13160301 | BAI35 | 72h | 31,802,103 | 76.70% |

**Supplementary Table B.** Lists of putative TALE targets after *Xoo* PXO83 and *Xoc* BAI35 infection.

[ Available as separate file target_list.xlsx ]

**Supplementary Table C.** Sequence logos of the binding motifs considered for TFs up-regulated after *Xoo* PXO83 infection.

| TF | motif ID | sequence logo |
|---|---|---|
| LOC_Os01g46800.1 | M07399_2.00 | 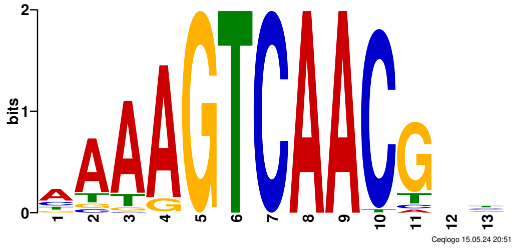 |
| LOC_Os01g61080.1 | M07391_2.00 | 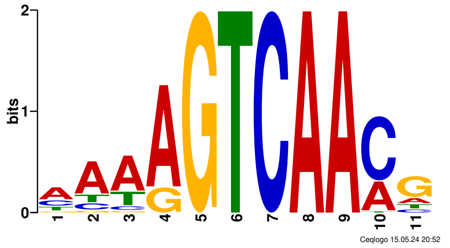 |
| LOC_Os01g63510.1 | M06905_2.00 | 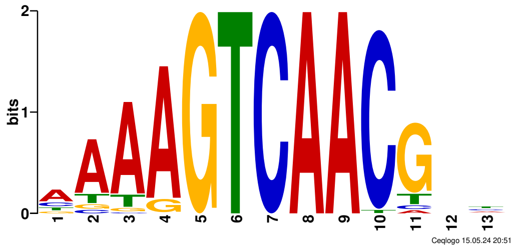 |
| LOC_Os09g29820.1 | AT5G38800.meme | 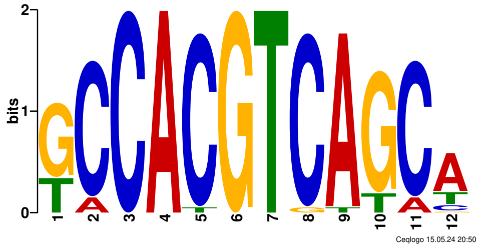 |
| LOC_Os09g29820.1 | Dimont/ChIP-seq | 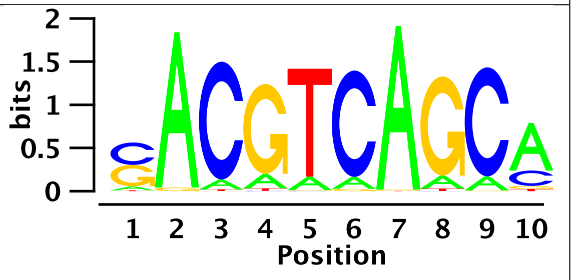 |

**Supplementary Table D.** Count values from read-based quantification, normalized expression values (DESeq2), time point-specific log2-fold changes and adjusted p-values for all rice transcripts according to the MSU7 annotation.

[ Available as separate file expression_data.xlsx ]

**Supplementary Table E.** List of RNA-seq data sets (ENA accessions) considered for generating co-expression networks

[ Available as separate file network_data.xlsx ]

**Supplementary Data S1.** RVD sequences and AnnoTALE names of TALEs present in *Xoo* PXO83

[ Available as separate file PXO83.fasta ]

**Supplementary Data S2.** RVD sequences and AnnoTALE names of TALEs present in *Xoc* BAI35

[ Available as separate file BAI35.fasta ]

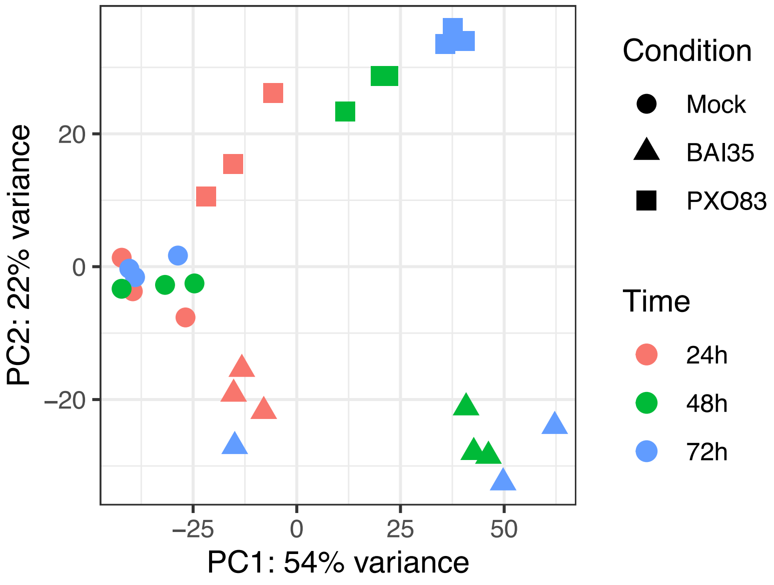


**Supplementary Figure S1.** PCA-plot of RNA-seq samples (vst-normalized expression values) after excluding all samples measured at the 36h time point.

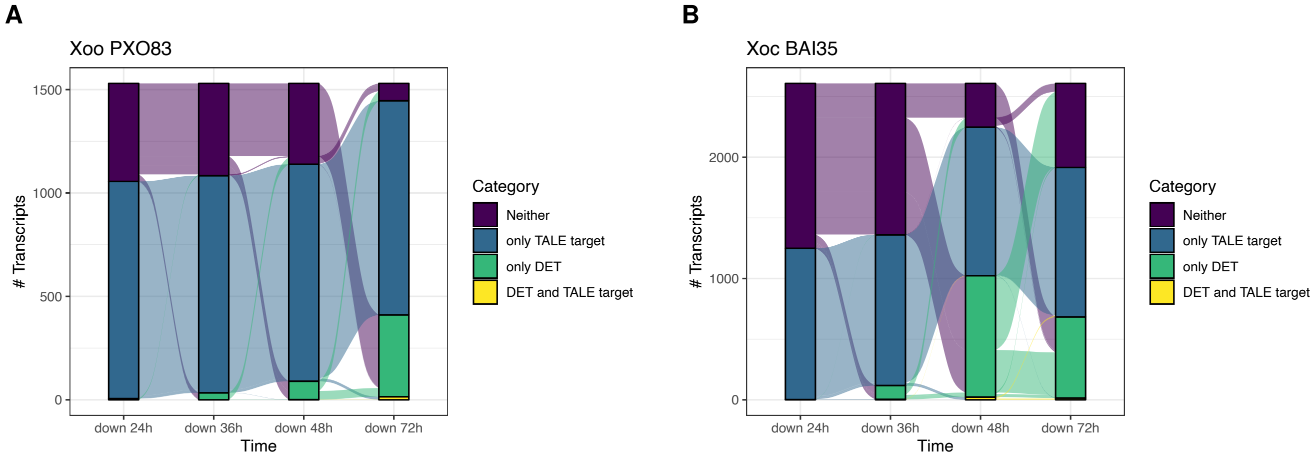


**Supplementary Figure S2.** Alluvial plot of down-regulated transcripts after *Xoo* PXO83 (A) and *Xoc* BAI35 (B) infection. Transcripts are classified in analogy to Figure 2.

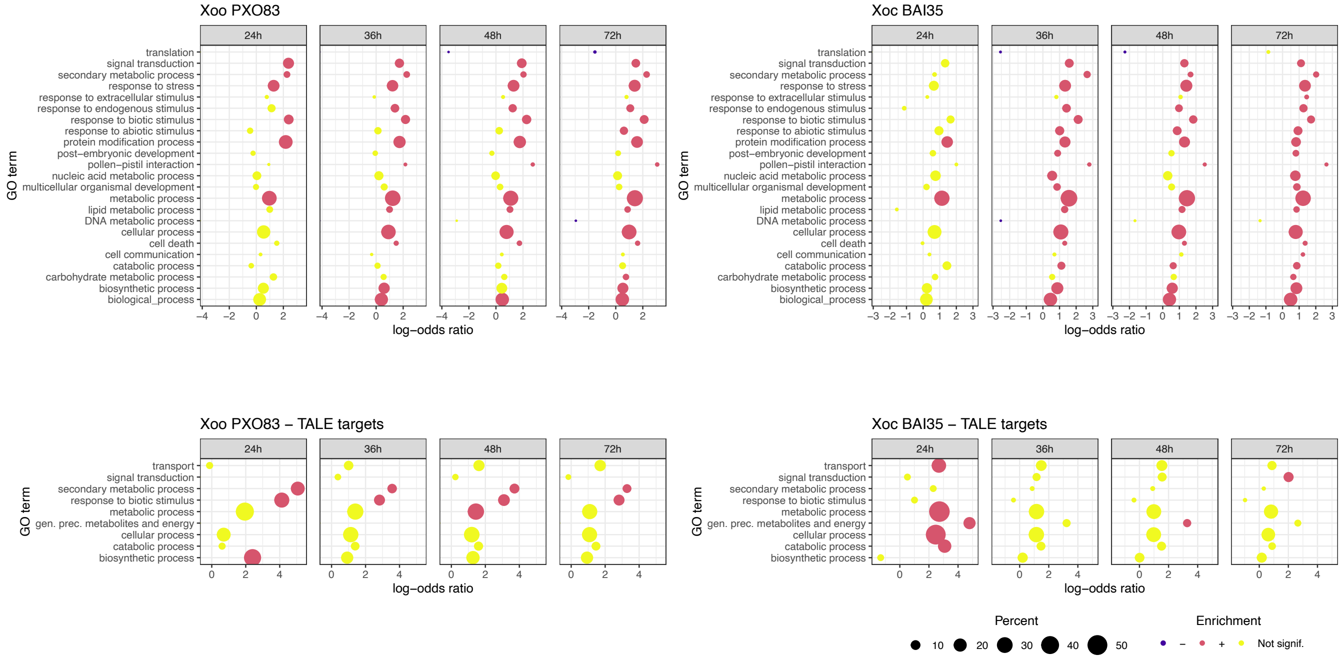


**Supplementary Figure S3.** Gene ontology (biological process) terms enriched among the DETs at each of the time points for *Xoo* PXO83 (left) and *Xoc* BAI35 (right) and separately for the respective TALE targets (bottom).

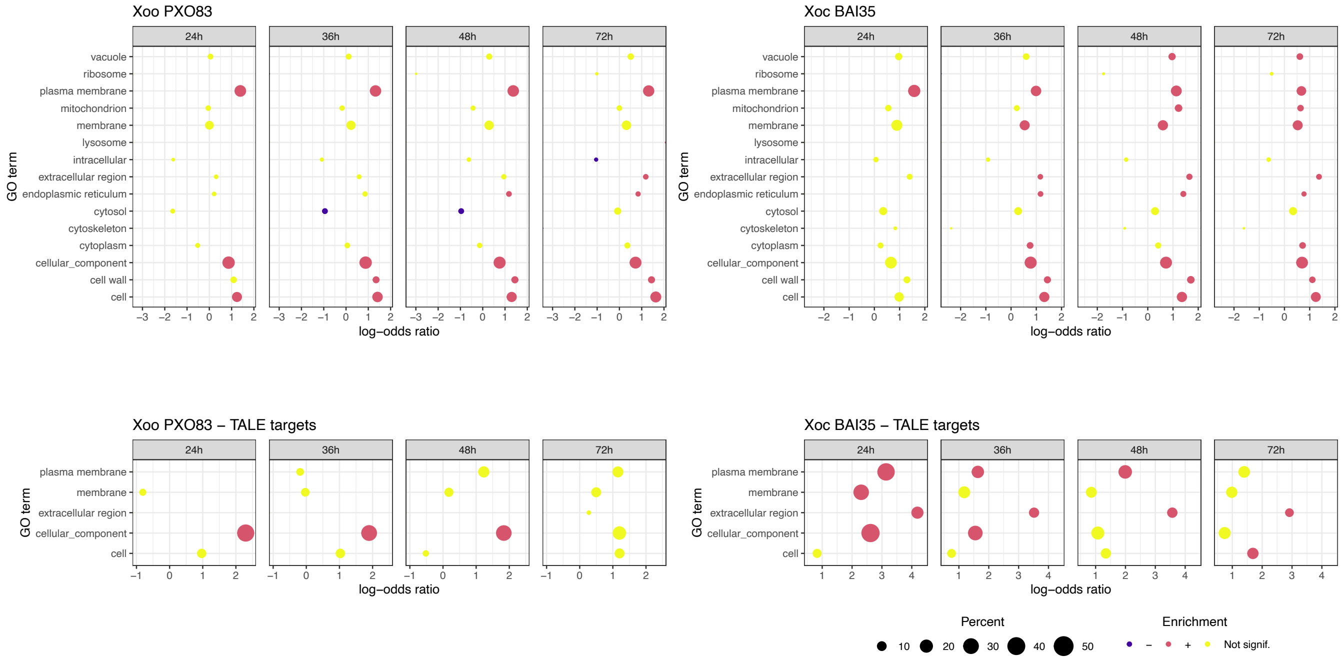


**Supplementary Figure S4.** Gene ontology (cellular component) terms enriched among the DETs at each of the time points for *Xoo* PXO83 (left) and *Xoc* BAI35 (right) and separately for the respective TALE targets (bottom).

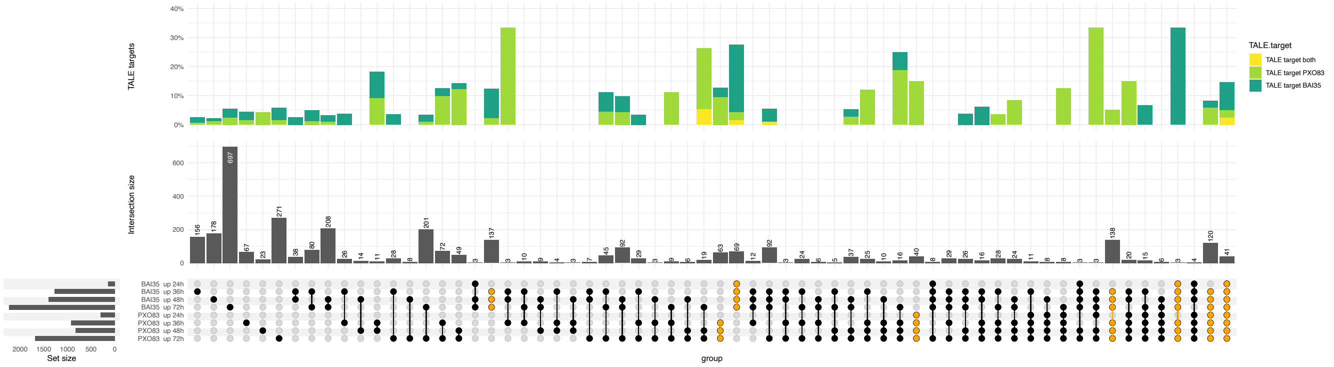


**Supplementary Figure S5.** Upset plot of all sets of differentially expressed, up-regulated transcripts after *Xoo* PXO83 and *Xoc* BAI35 infection. Subsets that cover contiguous time points within or across strains are highlighted in orange. The percentage of predicted TALE targets within each intersection set is indicated by additional barplots, where we distinguish between common TALE targets and TALE targets that are predicted for TALEs of only one of the strains.

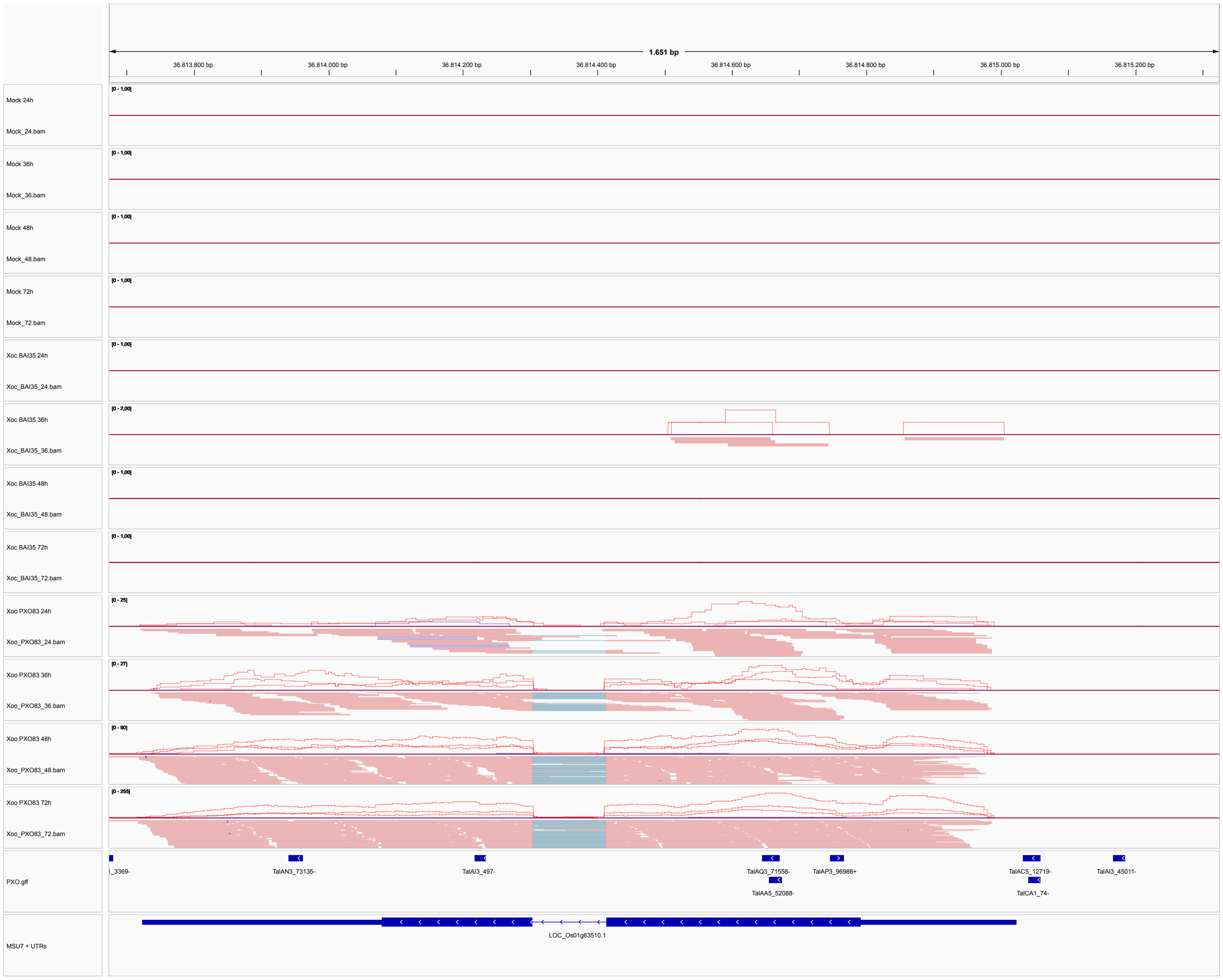


**Supplementary Figure S6.** IGV snapshot of an example transcript (LOC_Os01g63510.1) that is massively up-regulated after *Xoo* PXO83 infection and is predicted as a target of TalCA1.

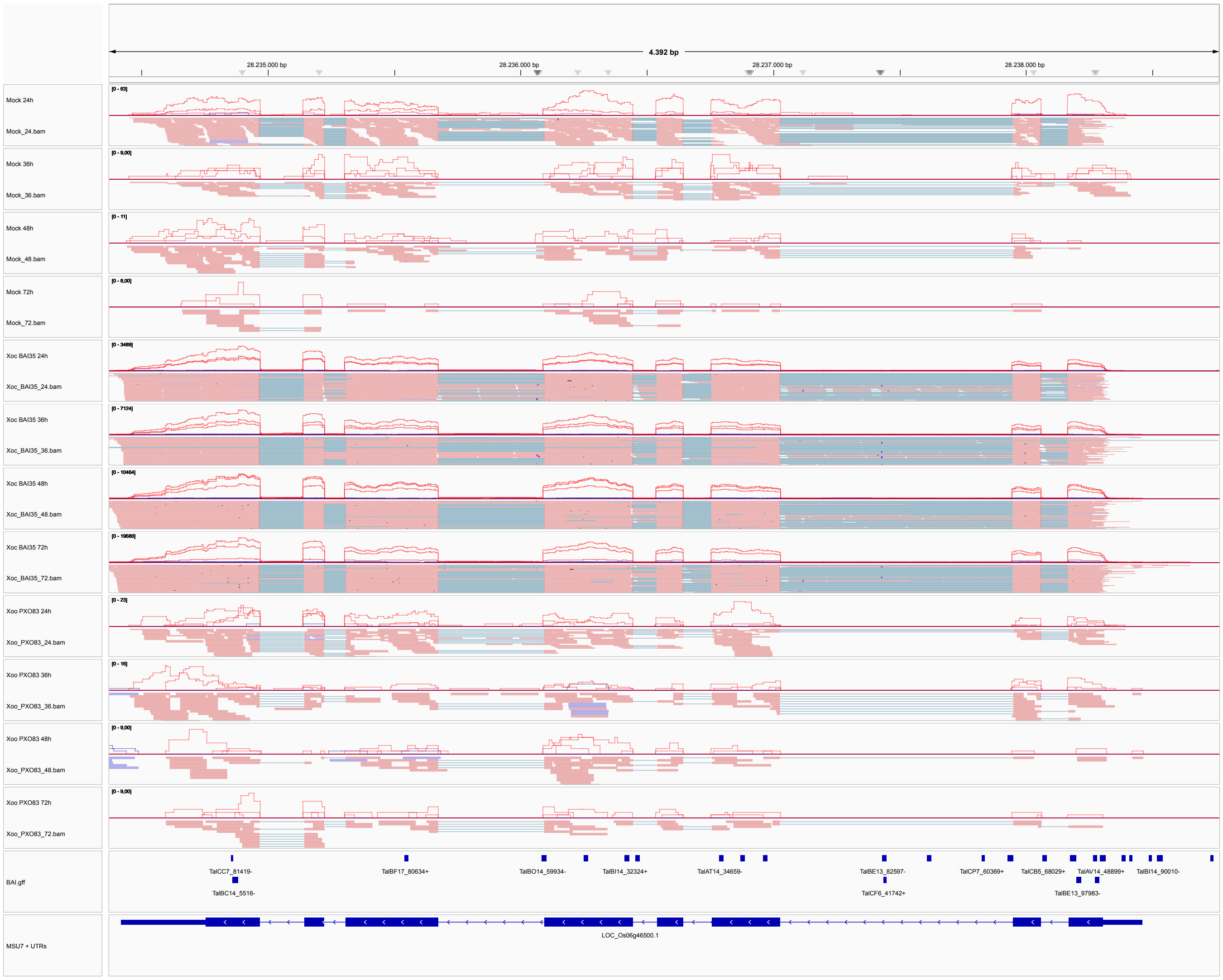


**Supplementary Figure S7.** IGV snapshot of an example transcript (LOC_Os06g46500.1) that is massively up-regulated after *Xoc* BAI35 infection and is predicted as a target of TalBF17.

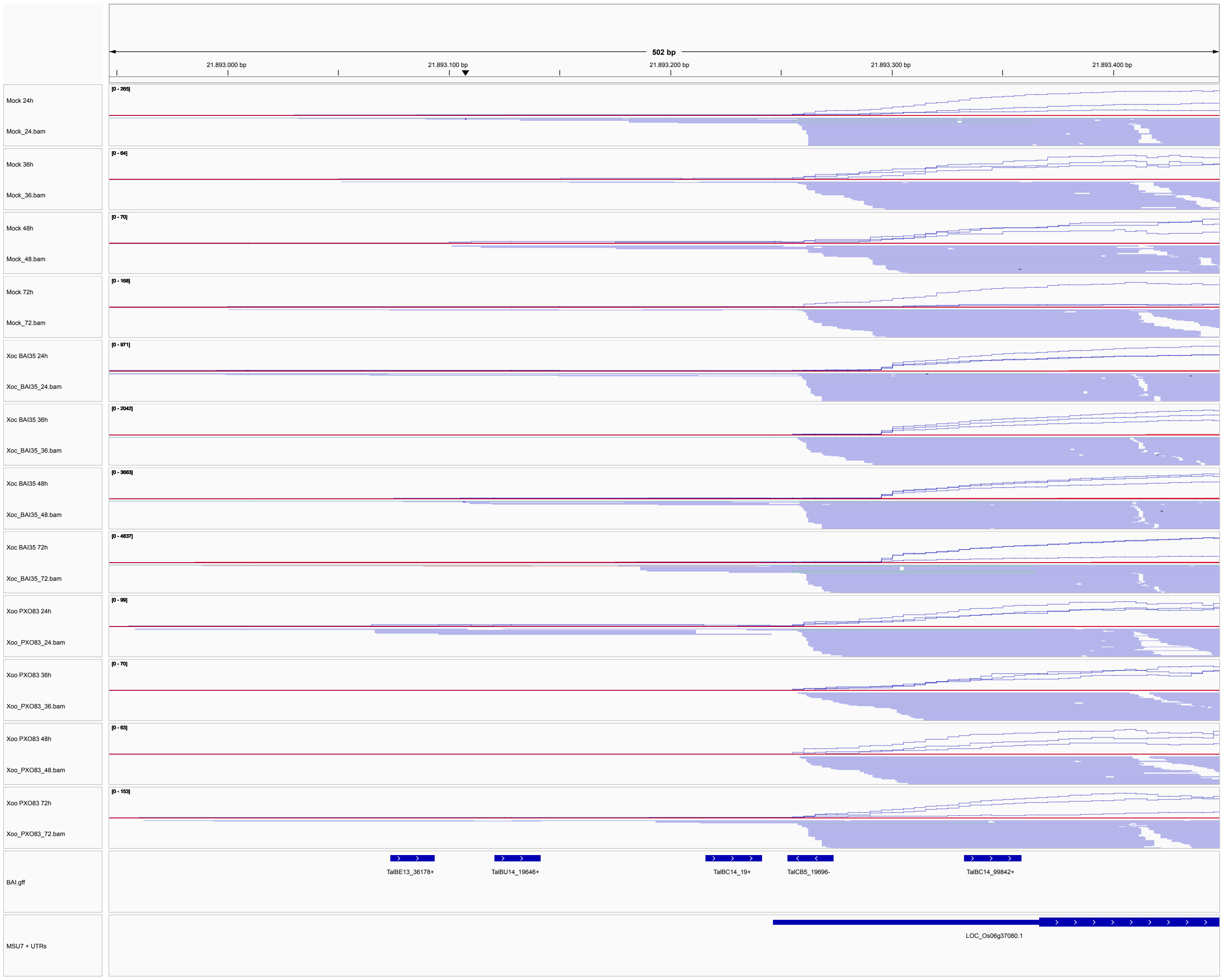


**Supplementary Figure S8.** IGV snapshot of an example transcript (LOC_Os06g37080.1) that shows up-regulation and a shifted transcription start site after *Xoc* BAI35 infection. While base-level expression continues from the natural transcription start site, coverage plots show that the increased expression mainly starts from a downstream transcription start site. Expression after *Xoc* BAI35 infection is predicted to be induced by TalBC14 and the corresponding target box is predicted at rank 19 in genome-wide predictions using PrediTALE.

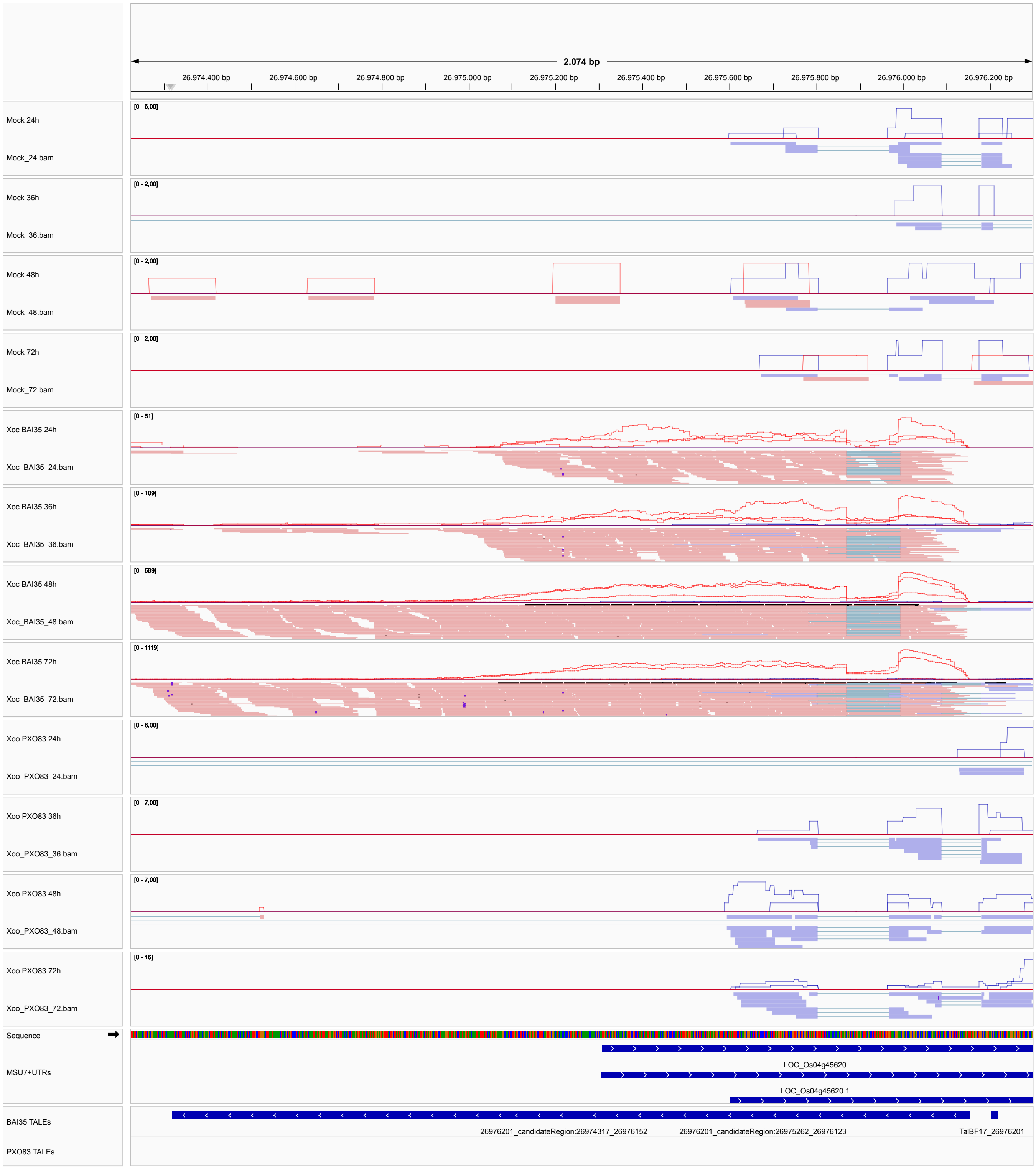


**Supplementary Figure S9.** IGV snapshot of an example transcript detected using DerTALEv2 based on genome-wide PrediTALE predictions that shows homology to a known gene from *Oryza glaberrima* (glycosyltransferase BC10-like).

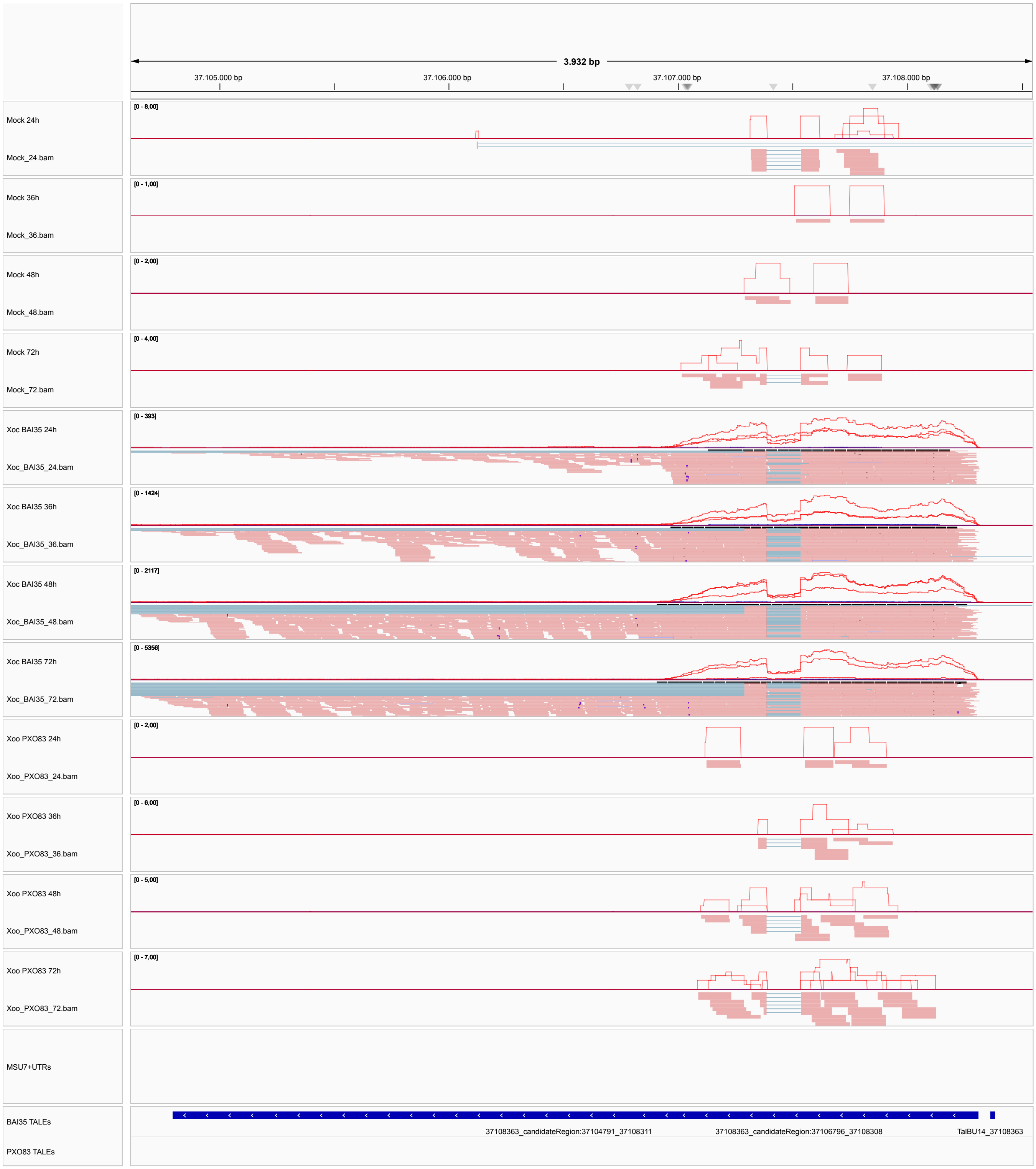


**Supplementary Figure S10.** IGV snapshot of an example transcript detected using DerTALEv2 based on genome-wide PrediTALE predictions that shows homology to a known non-coding RNA (SINE:OsSN1).

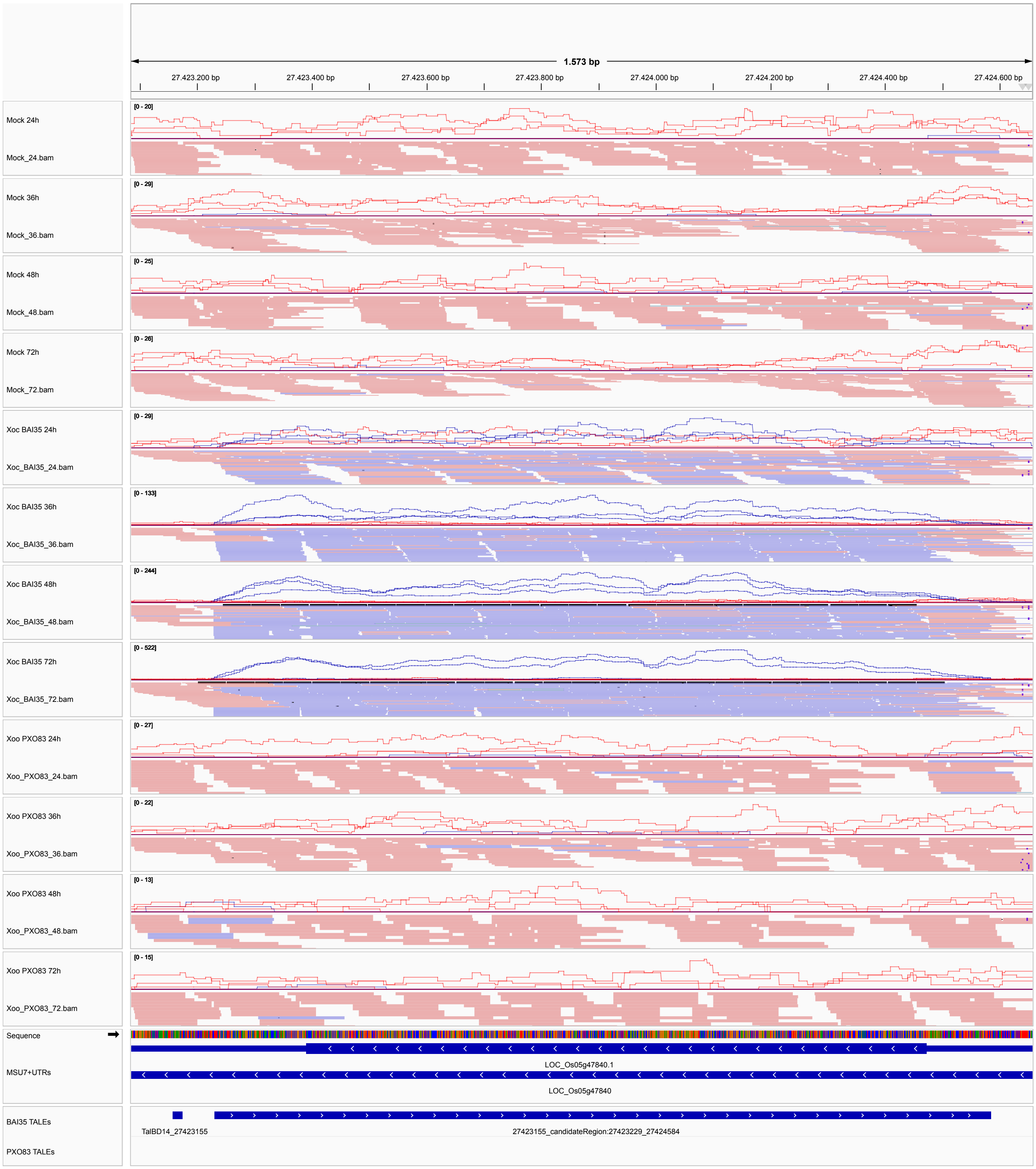


**Supplementary Figure S11.** IGV snapshot of an example antisense transcript (coverage and mapped reads on the forward strand; indicated by blue colour) detected using DerTALEv2 based on genome-wide PrediTALE predictions.

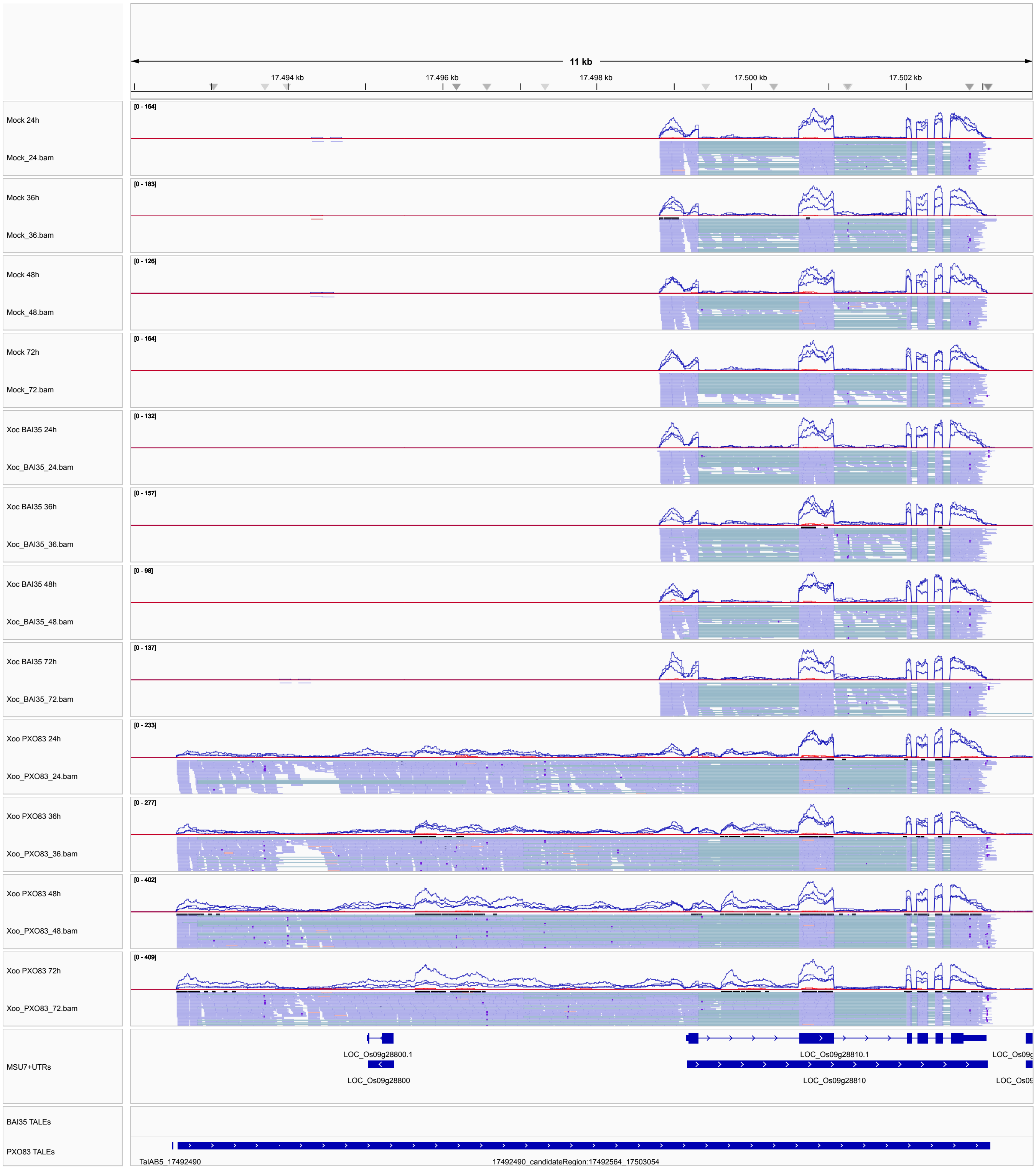


**Supplementary Figure S12.** IGV snapshot of an example transcript detected using DerTALEv2 based on genome-wide PrediTALE predictions that shows a massively shifted transcription start site and is predicted to be induced by TalAB5 after *Xoo* PXO83 infection.

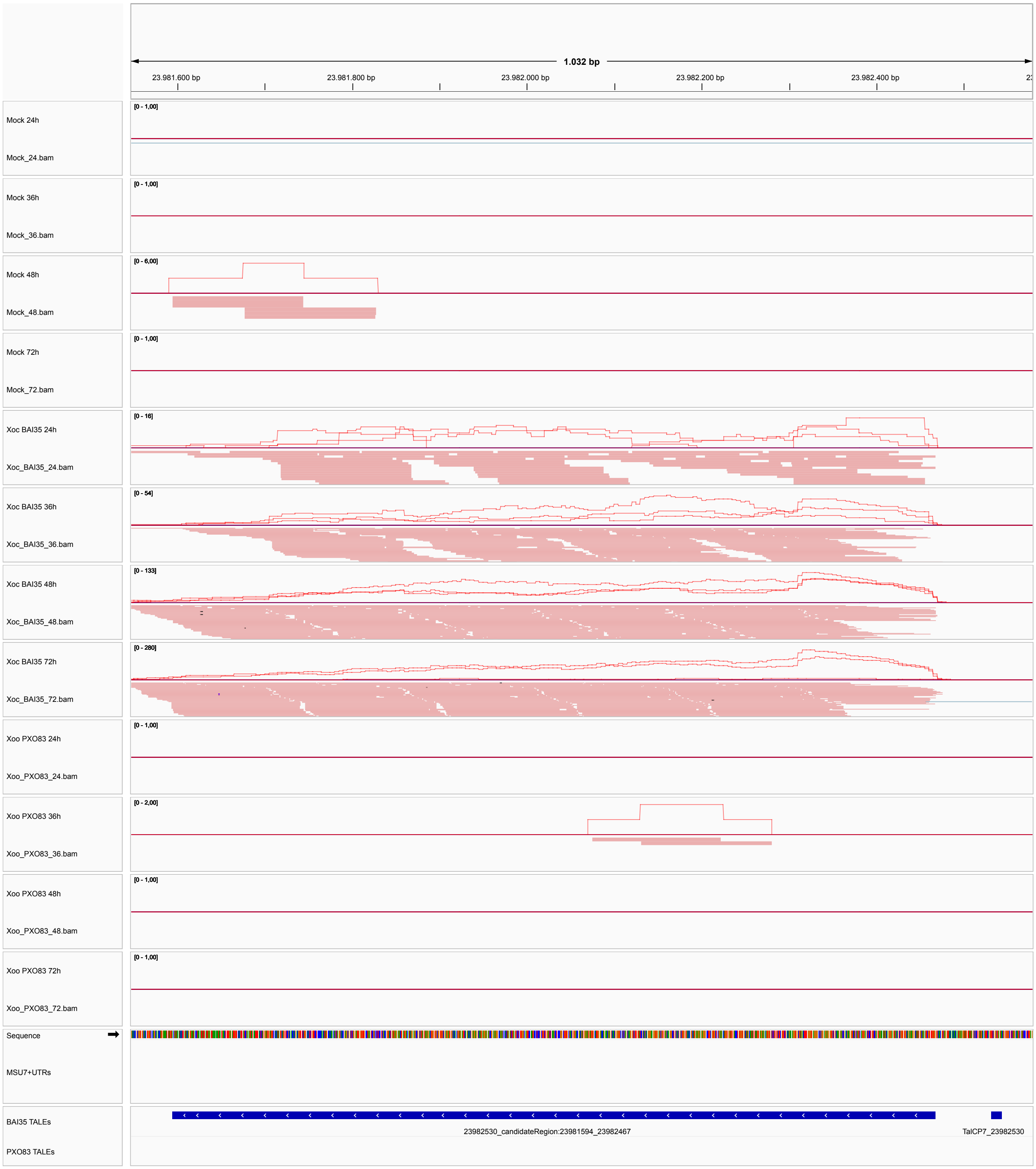


**Supplementary Figure S13.** IGV snapshot of an example transcript detected using DerTALEv2 based on genome-wide PrediTALE predictions without matches to known protein coding or non-coding transcripts according to NCBI Genbank, which is predicted to be induced by TalCP7 after *Xoc* BAI35 infection.

*Xoo* PXO83

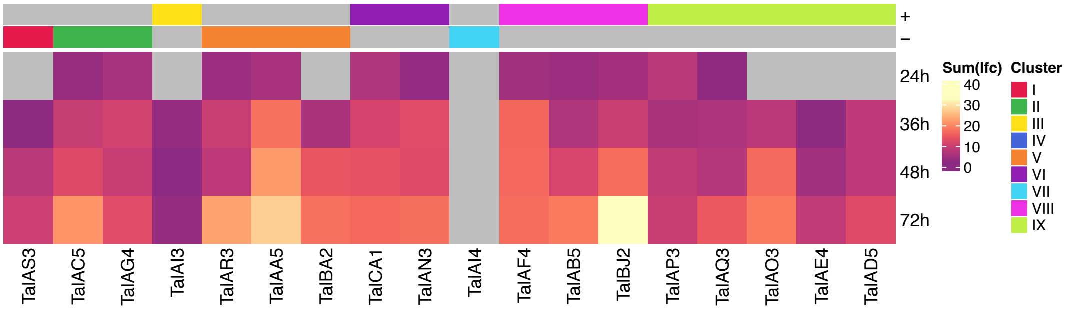


*Xoc* BAI35

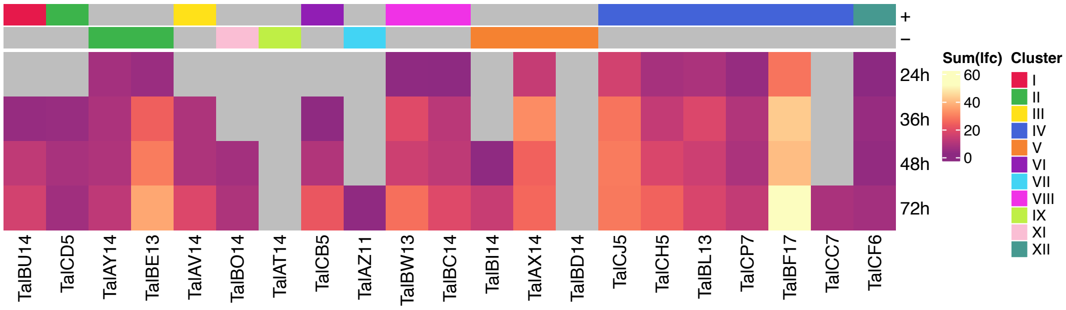


**Supplementary Figure S14.** Heatmap visualization of the sum of log2-fold changes of target genes per TALE after *Xoo* PXO83 (top) and *Xoc* BAI35 (bottom) infection. TALEs are ordered by their genomic organization and the respective TALE clusters and strand orientation are indicated by color annotations above the heatmaps. Here, a TALE target is counted if it is consistently up-regulated from the time point indicated at the ordinate until 72h.

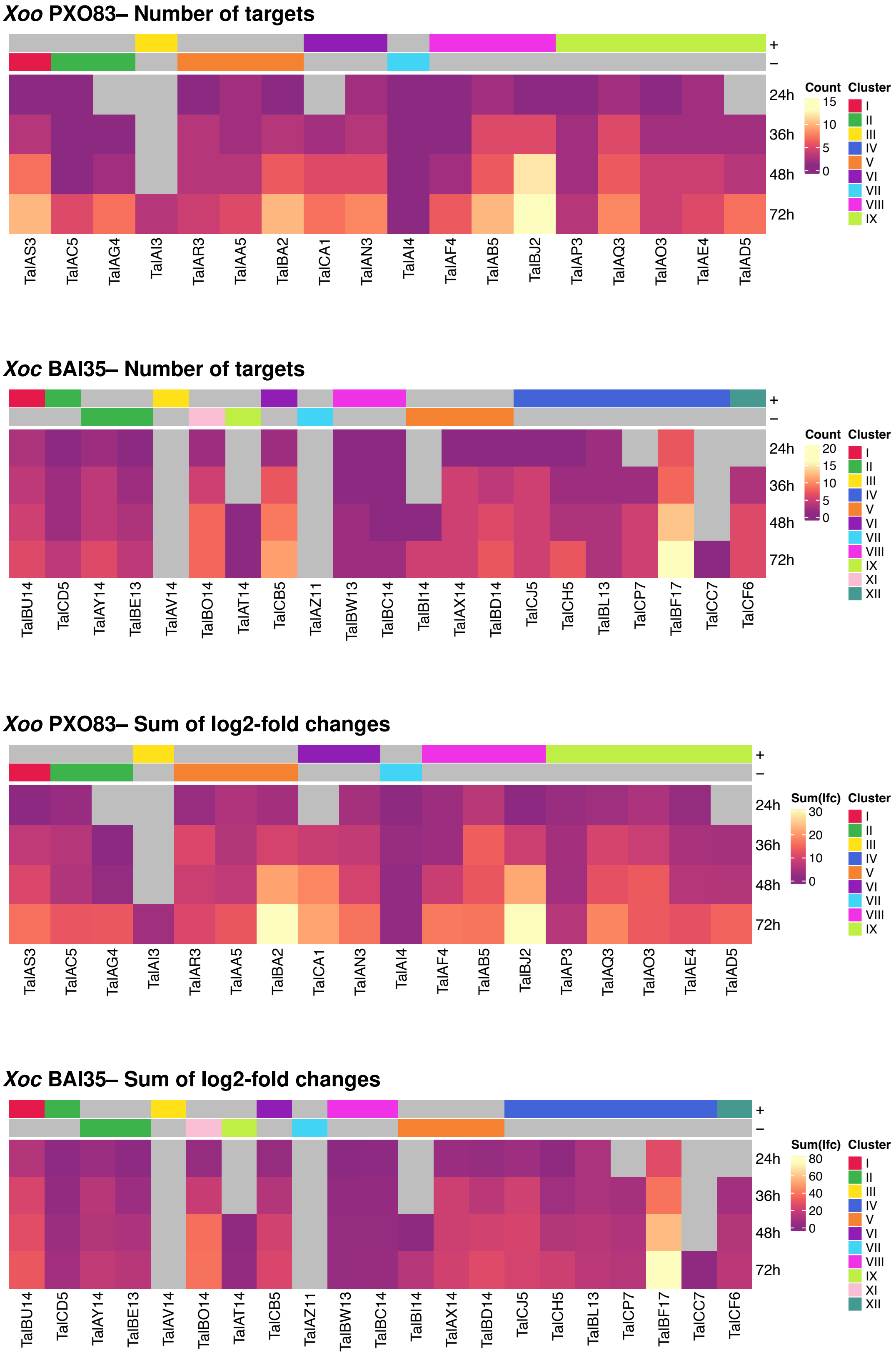


**Supplementary Figure S15.** Heatmap visualization of the number of predicted target genes (top) and sum of log2-fold changes of target genes (bottom) per TALE after *Xoo* PXO83 and *Xoc* BAI35 infection. TALEs are ordered by their genomic organization and the respective TALE clusters and strand orientation are indicated by color annotations above the heatmaps. Here, a TALE target is counted if it is consistently up-regulated from the time point indicated at the ordinate until 72h.

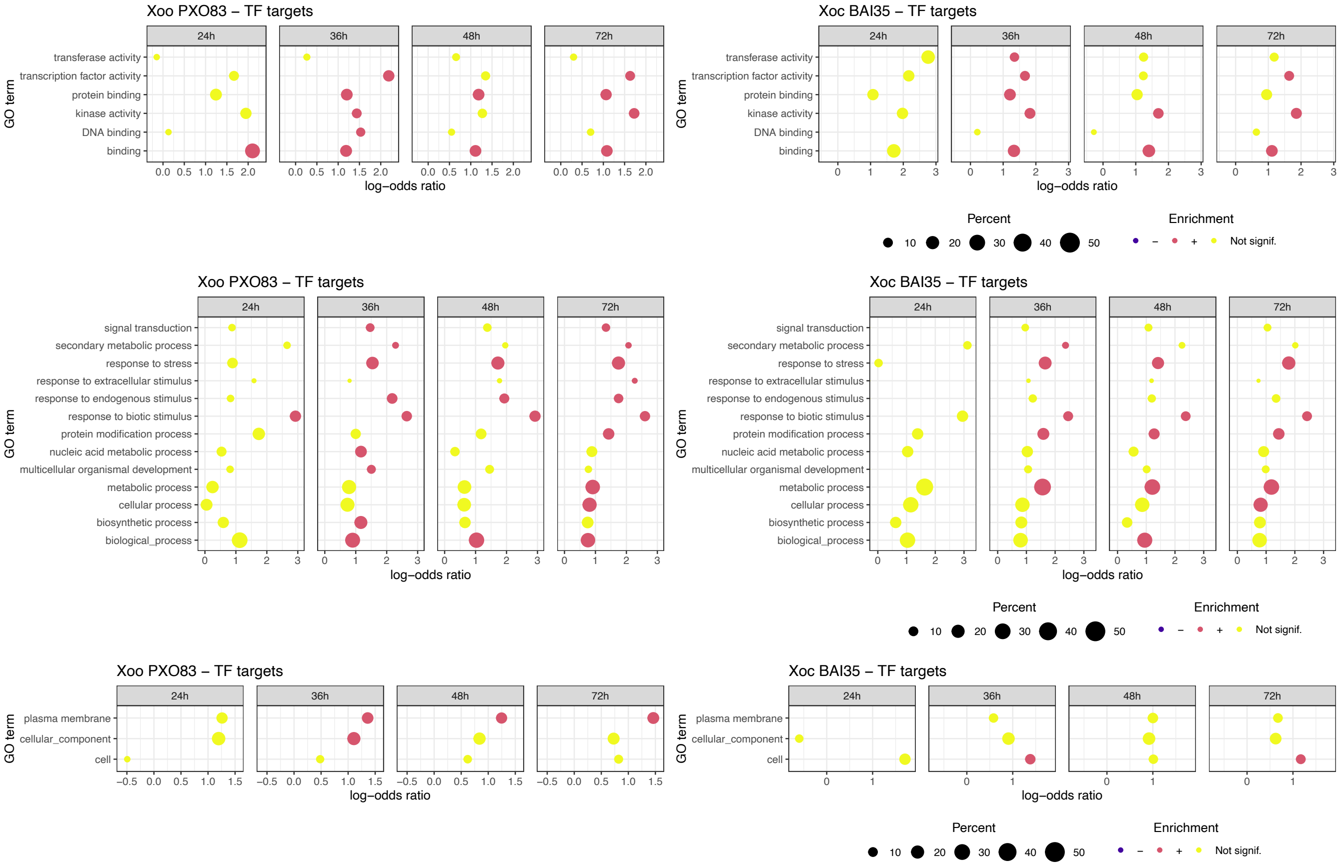


**Supplementary Figure S16.** GO terms enriched among the TF targets of TALE-induced TFs according to the "molecular function" (top), "biological process" (middle) and "cellular component" (bottom) ontologies.
